# Rolling 2D Bioelectronic film into 1D: a Suturable Long-term Implantable Soft Microfiber

**DOI:** 10.1101/2023.10.02.560392

**Authors:** Ruijie Xie, Qianhengyuan Yu, Dong Li, Xu Han, Xiaolong Xu, Jianping Huang, Wei Yan, Xiaomeng Zhou, Xinping Deng, Qiong Tian, Qingsong Li, Hanfei Li, Hairong Zheng, Guanglin Li, Fei Han, Tiantian Xu, Zhiyuan Liu

## Abstract

Long-term implantable bioelectronics can efficiently evaluate the function of the nervous system. 1D fiber-shaped devices require small surgical incisions significantly benefiting long-term biocompatibility. We report a strategy of rolling to reduce the dimension of bioelectronic devices from 2D film to 1D microfiber. The soft and stretchable multifunctional microfiber accommodates more than 60-channel longitudinally-distributed electrodes for bio-electric and -mechanical monitoring. It can be sutured into muscle with tiny incision for 8-month bio-electrical monitoring in rats. Importantly, after 13-month implantation, the fibroblast encapsulation around the fiber is negligible. Moreover, it can wander inside a brain under magnetic field. This strategy not only opens up the study of multi-channel and -function soft and stretchable fiber-shaped sensors but offers a platform for minimally-invasive implantable bioelectronics.

**One-Sentence Summary:** The rolling enables stretchable electrode array along one single microfiber and multimodal long-term *in-vivo* monitoring.

## Main Text

The central nervous and neuromuscular system provides the basic body control and kinetic energy for daily physiological activities of the human body, such as the movement of the body’s limbs (*1, 2*), heart stripes (*3, 4*), intestinal peristalsis (*5, 6*), and so on. During the operation, the neuromuscular system, for example, emits both electrophysiological (*7*) and mechanical signals (*8*) that can be monitored by sensors. Notably, soft sensors designed with mechanical properties similar to human tissues and the ability to stretch alongside them have proven particularly valuable in this context (*9-12*). These signals are vital for evaluating the functional state of the central nervous and neuromuscular system, and they hold significant implications for a wide range of applications, spanning human-machine interfaces to healthcare initiatives (*13-16*).

These soft and stretchable sensors are mainly categorized into body surface sensors and implantable sensors (*17-19*). Body surface sensors, attached to the skin, can only collect the average signal of the entire muscle bundles, particularly from the superficial muscles (*20-22*). Consequently, these sensors struggle to differentiate between various signals. In contrast, implantable sensor allows accurate monitoring of deep muscles and their individual components including muscle motor units (*23-25*). This implantable monitoring is desirable for precise understanding of muscles working process and significantly helpful for the control system in human-machine interfaces (*16*). For example, Max Ortiz-Catalan et al. implanted electrodes within the human body for up to 7 years to continuously collect myoelectric signals and control the prosthetic limb (*26, 27*). However, the rigid commercialized electrode, equipped with a few channels, provided limited control capabilities. So, how can we achieve long-term monitoring within the muscle bundle by more electrode channels to enrich the signal collection? Firstly, the electrode should be implanted in muscles with minimum surgery incision to minimize or eliminate the immune rejection effect, oppositely, larger incision induces severe blooding and releases more proteins attached to the electrode surface leading to severe fibroblast encapsulation (*28, 29*). It will largely increase the interfacial impedance of the electrode. In this regard, fiber-shaped electrodes could be a promising option, as they can be sutured into the muscle bundle with negligible damage to the surrounding tissue and muscles. In comparison, thin film electrodes necessitate complex and complicated surgical procedures, resulting in substantial damage to both skin and muscle tissue (*30*). Secondly, the electrode should possess soft and stretchable properties to match the characteristics of the target muscle (*31*). Thirdly, it should feature multiple channels along its length to effectively capture myoelectric signals throughout the muscle bundle. Therefore, a soft and stretchable fiber with multi-channel stretchable electrodes along its axis would be one of the most promising candidates for long-term implantable monitoring.

When considering fiber-like electrodes, the elongated thin film (hundreds of micrometers in width, several micrometers in thickness or thinner, and several centimeters in length) fabricated by the complicated photolithography has allowed for the longitudinal fabrication of multiple electrodes. However, the substrates used in this process are typically rigid and lack stretchablility (*32-35*). An alternative approach involves depositing the conductive film onto stretchable fibers through the dip-coating method. While it offers a simple system to investigate, it suffers from limitations in upscaling, the number of electrodes as well as delicate balance between electrode conductivity and stretchability (*36-39*). Furthermore, although the preform-to-fiber thermal drawing technique is emerging as a powerful and highly scalable approach to fabricate fiber-shaped sensors, the functionality is typically constrained at the fiber tip (*40*). Hence, the pursuit for a facile strategy to the fabrication of a stretchable microfiber with multiple longitudinally distributed channels remains unresolved. This endeavor becomes especially daunting when dealing with microfibers of diameters in the hundreds of micrometers range. Addressing this issue such as applying photolithography to a single soft microfiber, achieving encapsulation for multiple channels, and integrating various functions along the longitudinal section of the microfiber adds to the formidable challenges at hand.

In this article, we report a novel and simple strategy of rolling up a patterned nano-thickness stretchable film into a single microfiber, capable of accommodating up to 60-channel electrodes along one single microfiber. Remarkably, this rolling-up process results in the self-encapsulation of the electrodes, eliminating complex post-process treatment. Inside the fiber, the overlapping of adjacent electrodes forms a capacitive stretchable strain sensor, that simultaneously detects both bio-electrical and mechanical signals. The microfiber maintains its stable conductivity even when stretched to around 100% of its original length. It can be sutured into the muscle bundle with an incision of less than 200 μm in width, enabling continuous monitoring of both myoelectric signals and mechanical deformation of the muscle for at least 8 months. Furthermore, our extensive experimentation reveals that after up to 13-month implantation within rats’ muscle bundles, the inflammation effects and the fibroblast encapsulation (with only ∼23 μm in thickness) are significantly less pronounced as opposed to the control group implanted with the rigid fiber. These findings provide compelling evidence of the superior performance of the soft and stretchable fiber electrode in long-term implantation in muscle bundle compared to the rigid counterparts. Additionally, we highlight the impressive abilities of the soft microfiber electrode array to “submerge” into and “explore” the brain, facilitated by the assistance of magnetic force. The proposed strategy not only opens up the study of multi-channel and -function soft and stretchable fiber-shaped sensors, particularly those with longitudinally distributed function units, but also establishes a minimally invasive implantation platform for biomedical and healthcare applications that require prolonged monitoring of physiological signals.

### Rolling for the stretchable microfiber with multi-function electrode array

The basic idea to prepare the stretchable electrode array along one single microfiber is that a patterned thin-film electrode array is fabricated on sub-micro-thickness self-healing polymer with designed shapes first. Then the thin-film electrode will be rolled up during which the electrode array is self-encapsulated with detection sites and wire bonding points properly exposed finally (Fig. 1a). This is a dimension reduction process of the electrode from 2D film to 1D fiber (Movie S1). Compared to the film, 1D fiber will induce much less surgery incision and be desirable for the long-term implantable monitoring (Fig. 1b). Further, it could also inspire the fabrication of microfiber electronics with more complicated components inside following the same protocol.

**Fig. 1.**
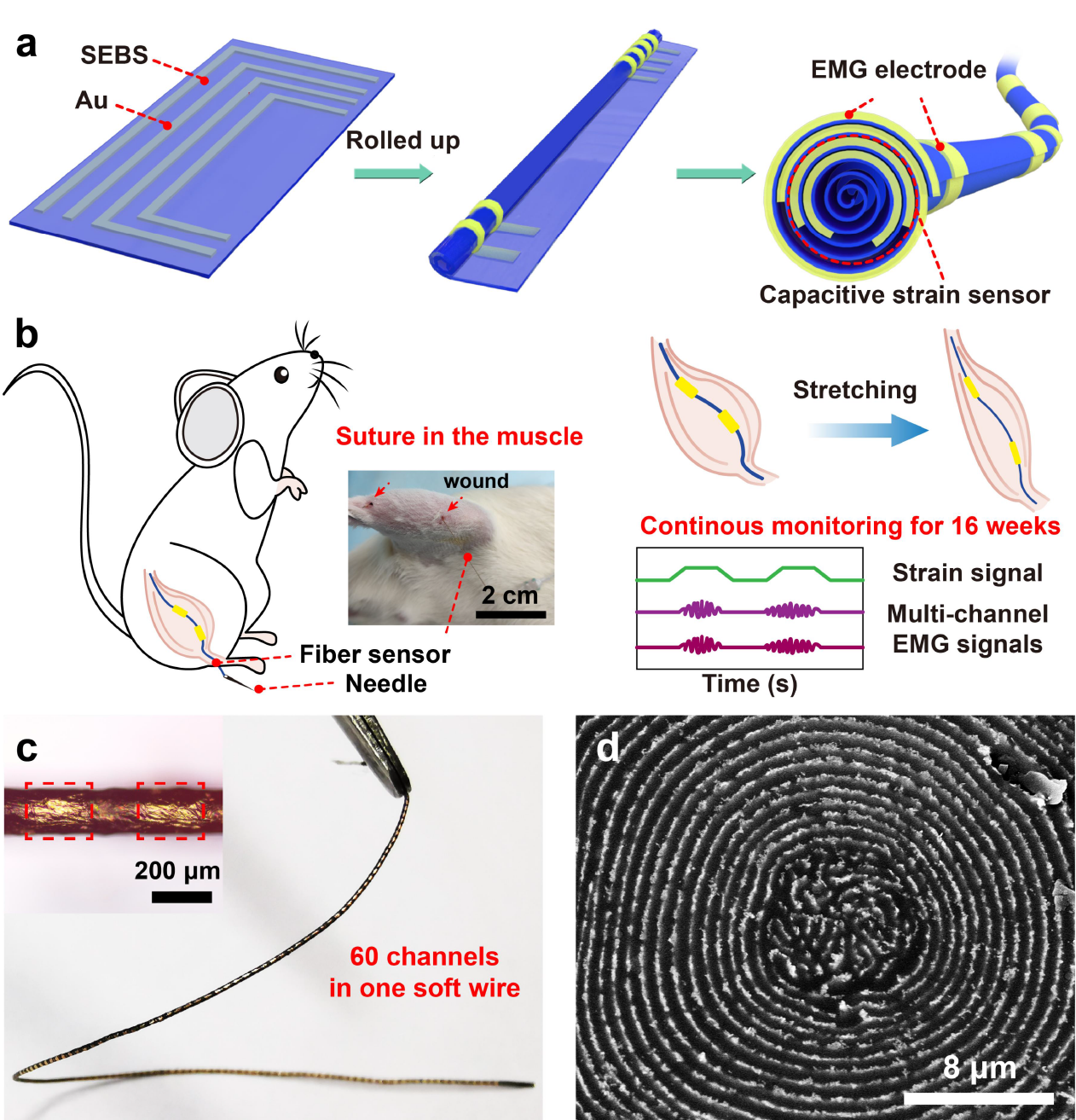
Preparation strategy and advantages of AoF-MSEA. a) Schematic diagram of the multifunctional fiber the preparation process. b) Fiber devices are sutured into rat muscle with a puncture needle and monitoring multichannel EMG and strain signals for 16 weeks. c) A fiber with 60-channel EMG electrode. d) SEM image of the cross section of the fiber sensor.

In detail, the conductive pathway on the film electrode was fabricated by depositing gold onto a 400 nm thick styrene ethylene butylene styrene (SEBS) film using vacuum thermal evaporation. The pattern of the conductive path is designed in both “C” shape and “L” shape. For the C-shaped pattern, when rolled up, two kinds of conductive points are exposed at both ends of the fiber: one for the electromyography (EMG) detection site and the other for wire bonding. In contrast, after two adjacent L-shaped patterns are rolled up, one side forms a in-plane capacitor, while the other side of these patterns serves as wire bonding points along the fiber (Fig. 1a).

The patterned thin-film electrode was first separated from the substrate and then rolled up to form a fiber. Due to the adhesion force between the thin film and substrate, an initial mechanical strain was induced in the thin-film electrode (Fig. S1a), which can also be confirmed from the SEM image of the fabricated fiber electrode (Fig. S1b). Thus, the successful fabrication of the fiber sensor requires consideration of two key factors. First, minimizing the adhesion force between the substrate and the film electrode is essential to streamline the roll-up process. In this regard, we chose polytetrafluoroethylene as the substrate due to its excellent hydrophobic properties, which significantly reduced the adhesion of the substrate to SEBS. Second, good stretchability of the gold film is necessary for the conductivity and stretchability of the final fiber electrode array. Thus, the stretchability of the composite film was investigated by varying the thickness of the gold films and the width of the conductor (Fig. S2a). Finally, we for the first time successfully crafted an along-one-single-fiber multi-function stretchable electrode array (AoF-MSEA) with more than 60 channels and the ability to detect both bio-electrical and - mechanical signals (Fig. 1c).

In addition, the diameter of AoF-MSEA varies depending on the width of the initial thin film. At present, it would be larger than 8 μm because the thin-film electrode was rolled up by hand and it will form an initial distorted layer inside at the very beginning of rolling followed by the normal rolling process (Fig. 1d). To improve the yield of AoF-MSEA by hand, a crucial step involves leaving an empty space of 1 cm in front of the conductive path (Fig. S4). Consequently, the minimum achievable diameter for the fiber is about 100 μm. Additionally, each additional 1 cm increment in the initial thin-film width results in an approximate increase of 24 μm in the fiber’s diameter (Fig. S5).

### Performance of AoF-MSEA

We employed a 3 cm wide thin film with four-channel stretchable conductors featuring two L and two C shapes to demonstrate the performance of AoF-MSEA. Each channel had a width of 2 mm with a 2 mm space between them (Fig. S4b). The diameter of the fabricated microfiber is ∼170 μm with proper wire bonding connecting to the hardware system (Fig. S6). Since the muscle fibers are stretchable during movement, AoF-MSEA should be also stretchable, thereby enhancing signal quality and reducing the risk of immune rejection. To assess the stretchability of the entire system, including the microfiber electrode array and its wire bonding part, we tested the conductivity when stretching the whole system (Fig. 2a). It revealed that the electrode array maintained excellent conductivity when stretched to 60% (Fig. S7). Interestingly, after 200 stretching cycles of 60%, its stretchability increased to 94% due to the rearrangement of micro cracks in the gold film (Fig. 2b). AoF-MSEA maintained low resistance after 1000 stretching cycles with 30% tensile strain applied (Fig. 2c).

**Fig. 2.**
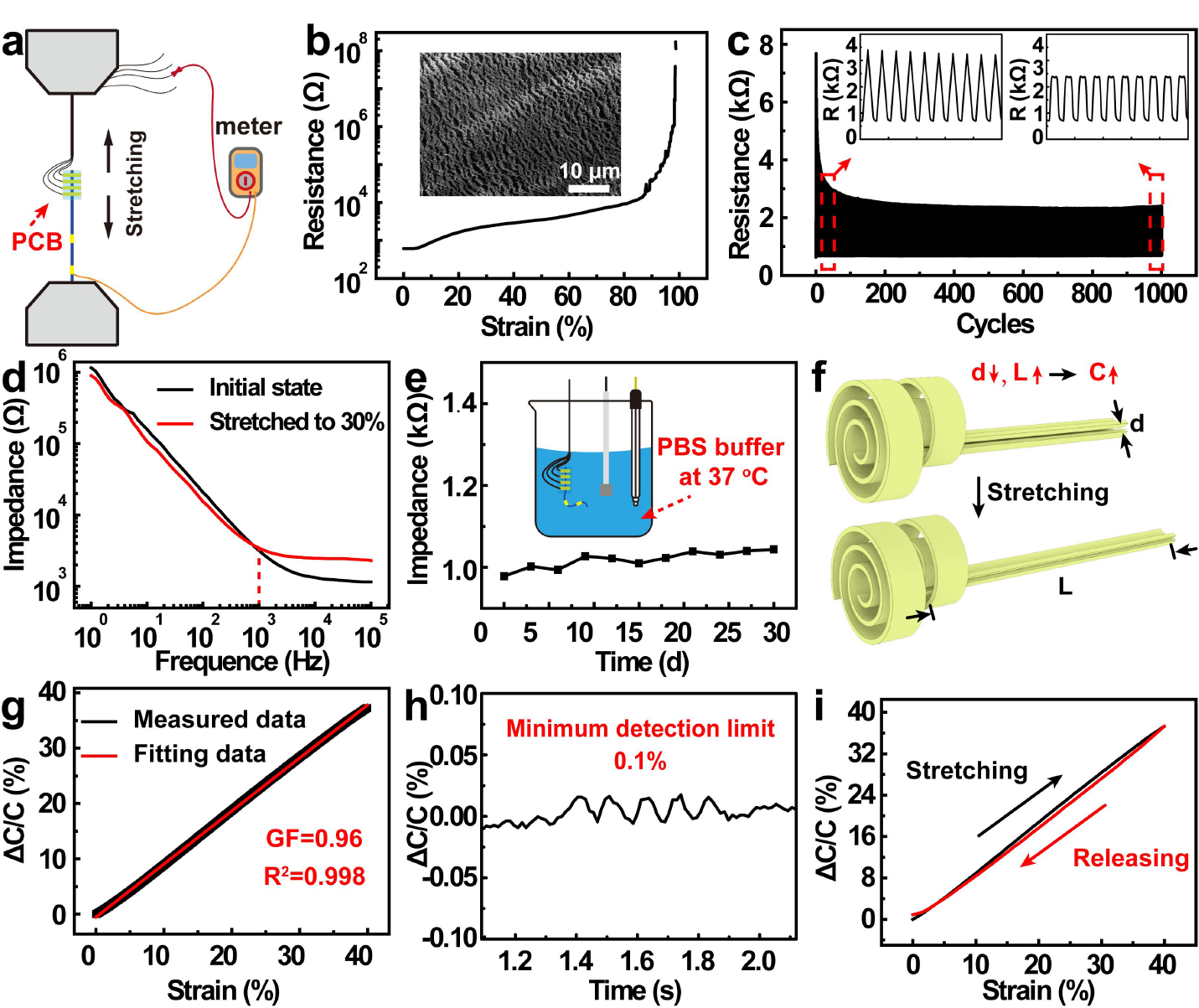
Performance of the AoF-MSEA. a) Illustration of the stretchability test for fiber sensor systems. b) Stretchability of the fiber sensor system, the image inside is the SEM image of the fiber sensor in the stretched state. c) Durability of the fiber sensor system after being stretched to 30% for 1000 cycles. d) Schematic illustration of the structural changes of the capacitor inside the fiber during stretching e) Change in the relative change of capacitance under various strains; GF was obtained by linear fitting. f) Hysteresis curve for fiber devices. g) Minimum detection limit of the fiber when used as capacitive stain sensor. h) Change in impedance of the fiber electrode when stretched to 30%. i) Waterproofing tests for the fiber sensor system.

Since the impedance between AoF-MSEA and tissue during movement significantly influenced the EMG detection, we examined the impedance of AoF-MSEA in phosphate-buffered saline (PBS) solution *in-vitro* first under different mechanical strain applied and *in-vivo* afterwards. When 30% of tensile strain was applied to the microfiber, the impedance slightly decreased at lower frequency range (e.g., < 1 kHz) and increased at high frequency range reflecting the increased resistance of the electrode. Since the targeted EMG signal is normally under 1 kHz, therefore, AoF-MSEA is supposed to stably detect the intramuscular EMG. Beside the interfacial detection site of the electrode, the wire bonding part of the system is also crucial for stable detection in implantable scenario. The permeation of interstitial fluid into the wire bonding interface can lead to the circuit failure between AoF-MSEA and PCB (See detailed structures in Fig. S6). To address this concern, we implemented two waterproof layers of para film and 703 silicone to enhance the waterproofness of the interface. Then, AoF-MSEA system was immersed in phosphate buffered saline at 37 °C to replicate its working environment *in vivo*. The impedance of AoF-MSEA showed negligible variation over the course of 30 days. This observation underscored the potential of the AoF-MSEA system for long-term implantable monitoring.

Moreover, the in-plane capacitor formed by two L-shaped channels underwent deformation with the stretching of AoF-MSEA. This enabled the fiber to be a stretchable capacitive strain sensor, monitoring the deformation of the muscle meanwhile (Fig. 2d). The fiber sensor exhibited a gauge factor of 0.96 and demonstrated excellent linearity (R-square value 0.998) within a strain range of 40% (Fig. 2e). Notably, as the fiber underwent stretching, the SEBS thin film between the two channels became thinner. It was found that when it was stretched beyond 40%, it will suffer electrical breakdown indicating its detection range. It was proved that the fiber sensor can detect minute mechanical strain as surprisingly low as 0.1 % (Fig. 2f), making it suitable for capturing the subtle muscle deformation. It was also observed that there was no obvious hysteresis for the stretchable capacitive strain sensor (Fig. 2g) with good repeatability (Fig. S8). In addition, our approach is not restricted to the SEBS substrate, but is also applicable to other polymer materials, e.g., ethylene-vinyl acetate copolymer or polydimethylsiloxane (Fig. S9). It is worth emphasizing that the good tear resistance of the film largely benefits the outcome of AoF-MSEA. Polymer films with low tear resistance are prone to fail in AoF-MSEA production due to the mechanical strain induced in the rolling process. This indicates that following our strategy, different materials and device components could be fabricated in 2D film first and then reduce dimension to 1D fiber by rolling.

### Wandering in the brain of AoF-MSEA guided by magnetic field

Brain-computer interface is becoming more practical in terms of algorithm especially with the assistance of AI big model, while the electrode array to extract information from brain still requires to be optimized. For example, how to implant the electrode into brain? The implantation method of the electrode determines the incision size and the ability to reach detection target in brain. Here, we demonstrate that AoF-MSEA can wander like an earthworm in brain to a specific detection site monitoring bio-electrical signals. This was achieved by mounting a magnetic head at the tip of AoF-MSEA, then the fiber can be guided by the external magnetic field.

In detail, since brain is enclosed within the skull, existing implantation methods often necessitate skull opening, causing big damage to patients and posing risks to their recovery. Based our concept, AoF-MSEA was guided by magnetic field and moved into brain through a 1 mm diameter skull opening in rat (Fig. 3a). Then stereoelectroencephalography (SEEG) signals were successfully monitored (Fig. 3b and Fig. S10). Besides, during the implantation in brain, it is important for the electrode to avoid the blood vessel. To demonstrate the dexterity of our strategy, we utilized a hydrogel-filled cerebrovascular scaffold to emulate a brain (Fig. S11). AoF-MSEA with the magnetic head wandered in the brain controlled by an external rotating magnet (Fig. 3c and 3d). The position and orientation of AoF-MSEA was controlled by tuning the position and rotation speed of the magnet. It facilitated three-dimensional maneuverability of the fiber electrode in brain, enabling it to avoid from puncturing blood vessels and reaching specific positions in brain (Movie S2). Furthermore, to simulate how AoF-MSEA collects signals during movement, we arranged five pieces of conductive hydrogel side by side, each applied with a different signal source (Fig. 3e). As AoF-MSEA passed sequentially through them guided by a magnetic field, related analog signals were detected at various locations (Fig. 3f). These outcomes suggest that we propose a new implantation method of electrode in brain by utilizing 1D fiber electrode array. Following our strategy, more active components, like tiny cameras, could be integrated in this fiber based on rolling. This will open up a new way to construct the human-computer interface with superior properties of multi functions, minimum implantation incision, and high maneuverability in brain.

**Fig. 3.**
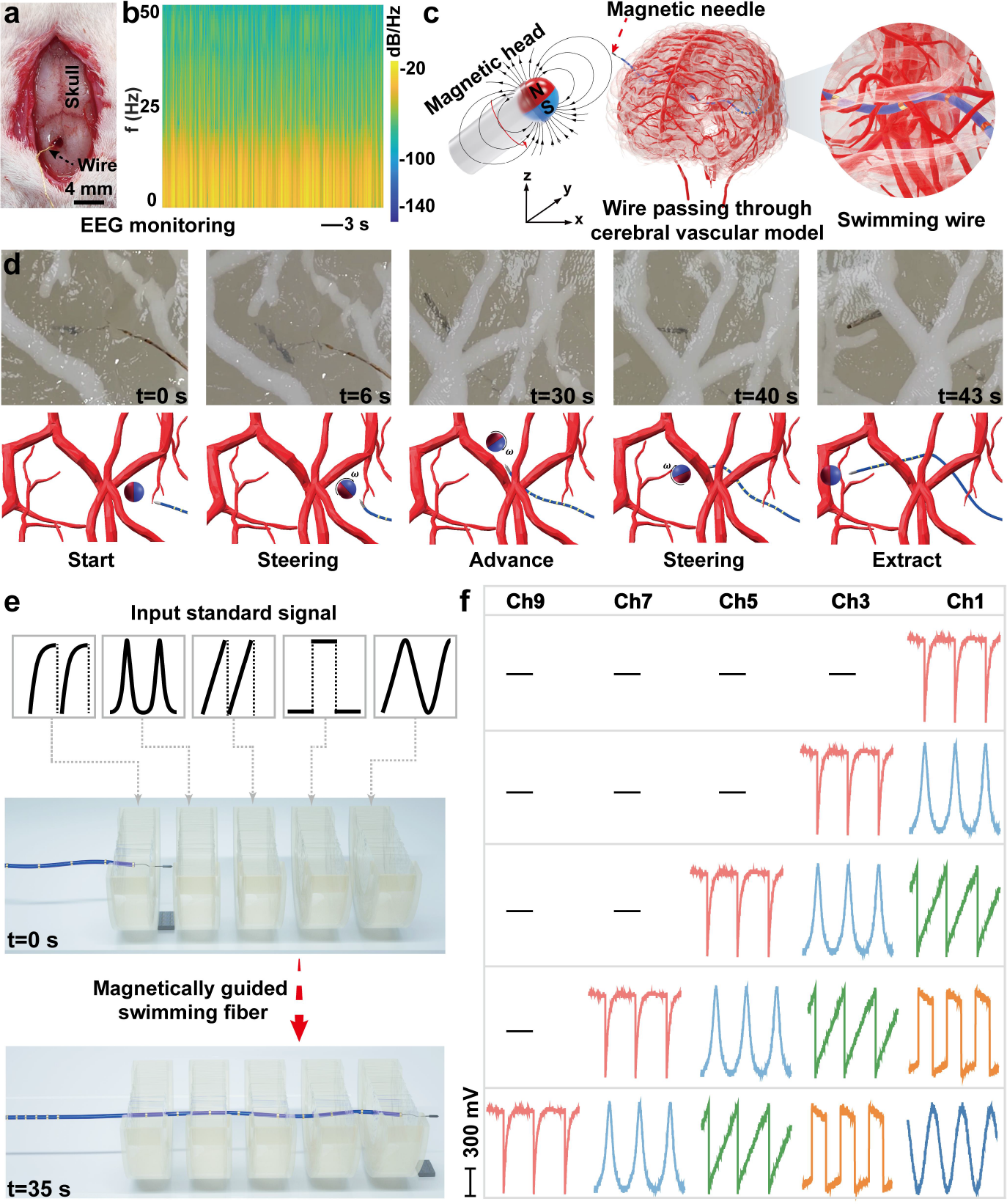
Implantation of AoF-MSEA under magnetic field. a) Implantation of fiber electrodes into the rat brain through a small aperture. b) Continuous monitoring of electroencephalogram (EEG) frequency spectrums within 10 min through the AoF-MSEA. c) Schematic illustration of swimming fiber in 3D printed cerebral vascular model guided by rotating magnetic head. d) The images and schematic diagrams of controllable swimming in agar-infused cerebral vascular model of fiber electrode in 43 seconds. e) Schematics of swimming fiber passing through five simulated neural units in sequence driven by external magnetic force. f) Recorded analog signals by swimming fiber from five channels (Electrode 1 (Ch1), Electrode 3 (Ch3), Electrode 5 (Ch5), Electrode 7 (Ch7), Electrode 9 (Ch9)) in the process illustrated in Fig. 3e.

## Long-term implantable monitoring of bio-electric and –mechanical signals in muscle

Peripheral neuromuscular system is the most important motion executor of human body. It is important to monitor its bio-electric and -mechanical signals *in-vivo* for healthcare and motion intention recognition. AoF-MSEA can be sewed into muscle just like a surgical suture with a minimum incision (Fig. 4a, S12 and Movie S3). The fiber was affixed to a needle and gently inserted into the muscle. The wire bonding part with connecting wires went under skin and mounted to an adapter as the female head fixed on the rat’s skull (Fig. 4a, 4b). Thus, the adapter was not easily bitten off or grabbed away by rats. After surgery, rats can walk freely. Signals were recorded via inserting a male head to the female one (Fig. 4c, 4d, Movie S4). Fig. 4e and 4f depict the top view and X-ray imaging of the electrode implantation pathway schematic. We optimized the whole setup for years to stabilize this system. Many small issues, including skull fixture manner, encapsulation of wire bonding, wire connection manner, etc., can make the signal collection fail, although AoF-MSEA itself is stable. We decide to release the whole setup information on a website (*41*). Now, it shows the whole frame and more details are being filled in. Hopefully, it could save other researchers’ time.

**Fig. 4.**
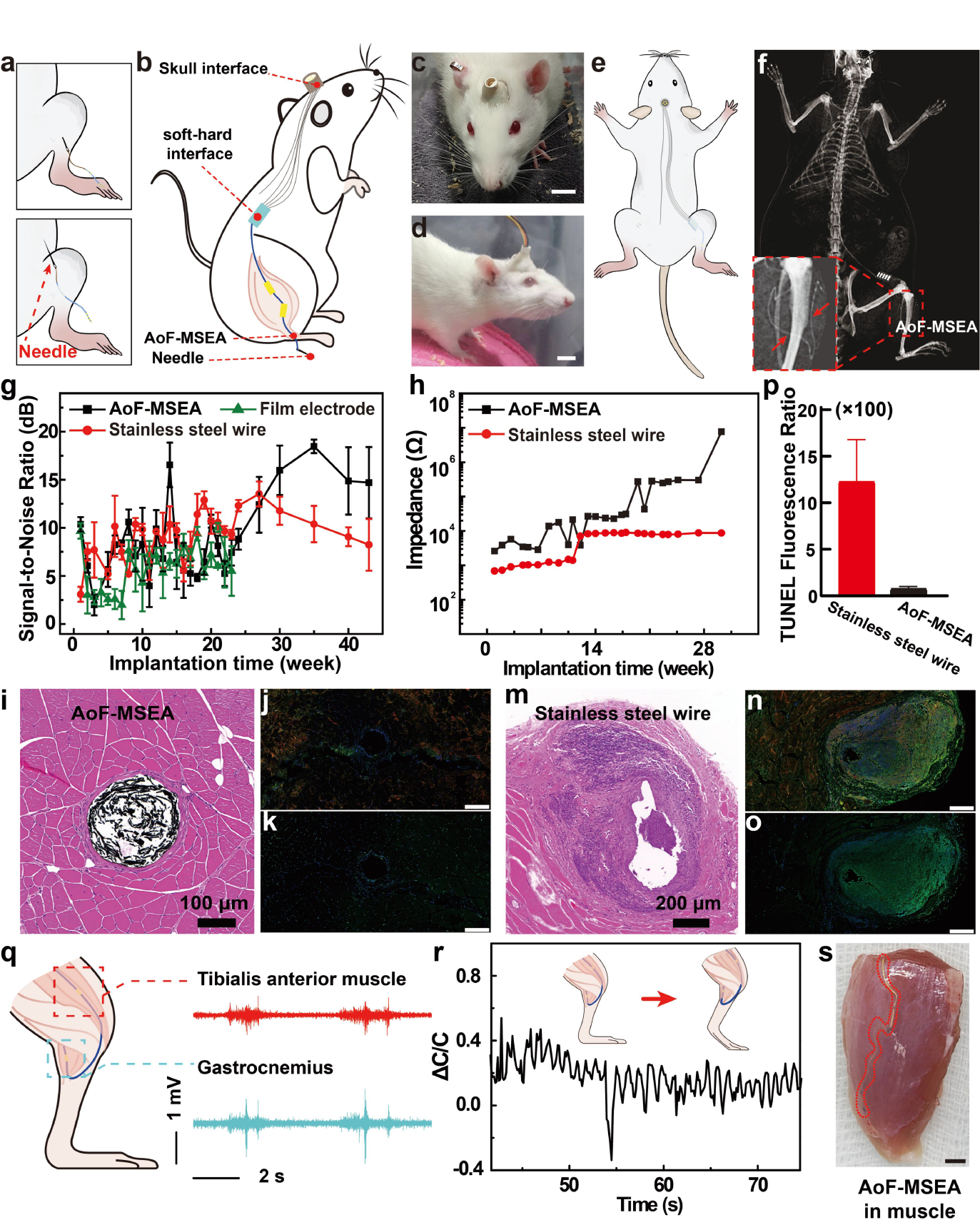
Long-term monitoring performance of fiber sensor for muscle information. a-b) Minimally invasive electrode implantation method and schematic diagram of the implantation pathway in rats after implantation. c-d) The images of skull interface and signal transmission interface. cale bar: 1 cm. e-f) The distribution diagram of implanted electrodes in the rat’s body, and the X-ray images of the electrodes. g) Comparison of signal-to-noise ratio of AoF-MSEA, stainless steel wire and thin film electrode over 43 weeks after implantation. h) The impedance trend for AoF-MSEA and stainless steel wire implanted for 30 weeks. i) Hematoxylin-eosin staining results of tissue around AoF-MSEA. j-k) Immunofluorescence staining result (j) and TDT-mediated dTUP Nick-End Labeling result (k) of tissue around AoF-MSEA, respectively. Scale bar: 200 um. m) Hematoxylin-eosin staining results of tissue around stainless-steel fiber sensors. n-o) Immunofluorescence staining result (n) and TDT-mediated dTUP Nick-End Labeling result (o) of tissue around the stainless steel wire, respectively. Scale bar: 200 um. p) TUNEL fluorescence ratio comparison between AoF-MSEA and stainless steel wire. (Calculations were conducted using image processing software, ImageJ. Statistical analyses were performed using GraphPad Prism. Significance was assessed through the Mann-Whitney test, with p-values less than 0.05 considered to have statistical differences.) q) Simultaneous acquisition of EMG signals from two muscles using the fiber sensor system. r) Muscle strain monitoring using a fiber sensor system when stretching the rat’s leg. s) Anatomy sample with AoF-MSEA embedded in the muscle. Scale bar: 1.5 mm.

Before long-term implantation of AoF-MSEA, acute trials were conducted to ensure the basic detection ability of the fiber electrode. Also, acute intramuscular EMG monitoring is a standard clinical method for assessing nerve damage. Currently, the used electrode is a rigid needle with diameter of around 300 μm. When muscles of patients contract as ordered by a doctor, the rigid needle agitates muscles evoking intense pain. So, soft and stretchable fiber-shape electrodes become the ideal substitute. To verify this, the intramuscular EMG signal induced by stimulation of the sciatic nerve was recorded by the fiber electrode (Fig. S13). As the applied current increased, the amplitude of the intramuscular EMG recorded increased accordingly (Fig. S13). The distinct channels displayed notable variations due to varying distances from the stimulation site, indicating that these channels do not interfere with each other during EMG monitoring. Then, the nerve situated between the muscle and the stimulation site was subjected to different degrees of clamping forces (Fig. S14). For a stimulation current of 36 μA, the amplitude of the EMG signals decreases with increasing levels of damage. In cases of severe nerve damage, even at a stimulation current of 200 μA, no intramuscular EMG was detected, highlighting the effectiveness of AoF-MSEA in assessing nerve damage.

The AoF-MSEA was subsequently implanted in rats for long-term monitoring of bio-electric and -mechanical signals from muscles. Long-term implantable monitoring is especially important for human-machine integration, e.g., prosthesis control and exoskeleton (*27, 28*). We implanted AoF-MSEA into the anterior tibial muscle and recorded the EMG weekly to evaluate its stability. The detailed description of electrode implantation experiments in animals, along with images as shown in Fig. S15-S21. For comparison, we also implanted normal 2D film-shape stretchable electrodes constructed by same materials and a rigid stainless-steel electrode with a diameter of 170 μm into the same muscle in the opposite leg of this rat. For 2D film electrodes, it induced a larger surgery incision with more blood released, hindering the contact of the electrode with the muscle and contaminating the electrode (Fig. S16 and S21). For the fiber-shaped stainless-steel electrode, it can be implanted in the muscle using the same method as AoF-MSEA, but it was rigid with high Young’s modulus of >10 GPa leading to tissue damage and long-term inflammatory response. It can not be stretched with the deformation of muscles. For our AoF-MSEA, since it was soft and stretchable, and when it was sutured into the muscle, it can keep contact with muscle offering a superior signal-to-noise ratio compared to the 2D film electrode and the stainless-steel electrode within 30 weeks (Fig. 4g). We tried this implantation of AoF-MSEA in five rats. One of them failed due to the occasional death of the rat. Four of them succeeded and the average time of long-term monitoring was around 10 months (40 weeks, Table S1). We also monitored the impedance change during those time finding that the impedance gradually increased and reached to around 1 MΩ after 28 weeks. Notably, the rigid stainless-steel electrode also kept low impedance in this 28 weeks and can detect intramuscular EMG continuously, although the inflammatory response was quite serve (Fig. 4g and 4h). This serve inflammatory may induce cancer later (*42, 43*). We attempted to analyze the reasons behind the above results. The signal-to-noise ratio data were derived from the analysis of EMG data during rat motion. As implantation time increased, the active space of the steel wire electrodes expanded. Combining this observation with staining data, it became evident that the stainless steel wire electrodes consistently induced a more pronounced inflammatory response in the surrounding tissue. The interaction of these factors led to a decreasing trend in the signal-to-noise ratio after an initial stable period. Conversely, the soft fiber electrodes, with tissue moduli similar to that of the surrounding tissue, evoked fewer rejection responses. As a result, their signal-to-noise ratio gradually increased after an initial stable period, although there was a decreasing trend in the later stages. And the rat impedance data were collected under static conditions and remained unaffected by interface changes during movement. Then we carefully analyzed the intramuscular EMG detected for a period of around 40 weeks (Fig. S22 and S23). Starting from the 24th week, the central frequency value was around 150 Hz, and then it decreased to around 50 Hz by the 43^rd^ week. The amplitude of the frequency spectrum became significantly lower after 24 weeks. These indicated that although the EMG was detected indeed, but part of the information lost due to the increased impedance of the electrode array. From the impedance change, it was observed that there was an impedance jump at around 20^th^ week and after 23^rd^ week, the impedance kept in high level (> 0.5 MΩ ohm).

After 13 months (54 weeks) implantation, we carefully conducted the taking out process of AoF-MSEA and rigid fiber electrodes (See details in Fig. S24-S26 and the website (*41*) (including the detail anatomical information for other four rats in Fig. S27-36)). After rats’ skin opening, it indicated that it was quite clean around the fiber electrode array by visual observation (Fig. S25d-e). However, there were several breaking points in the fiber probably due to the local deformation concentration (Fig. S24 and S25). Based on the impedance change (Fig. 4h), the breaking may occur at 21^st^ week when the impedance jump to high level. After that, the EMG signal should be detected by the conductor exposed in the fiber at the breaking point. To further analyze the bio-compatibility of different electrodes, we conducted the tissue hematoxylin-eosin (HE) staining in different rats respectively. For AoF-MSEA, it clearly showed that the thickness of the fibroblast encapsulation around AoF-MSEA was surprisingly small, less than 23 μm and did not induce any inflammation (Fig. 4i). We further observed the gold film surface in AoF-MSEA finding that although the fiborblast encapsulation was negligible, the surface was still contaminated with a layer of protein or other biomaterials (Fig. S32 and S33, compared with Fig. S37). This showed the evidence why the impedance gradually increased in the long term. In contrast, the thickness of fibrotic tissue around the stainless-steel electrode exceeded 451 μm (Fig. 4m). Furthermore, results from CD68-CTGF immunofluorescence double label staining and TdT-mediated dUTP nick-end labelling (TUNEL) indicated that the fluorescence intensity around AoF-MSEA showed negligible changes compared to normal tissue (Fig. 4j and 4k), while the stainless-steel electrode displayed a significant increase in fluorescence intensity (Fig. 4n and 4o). This suggested that compared to the rigid stainless-steel fiber electrode, AoF-MSEA possessed a much lower concentration of connective tissue growth factor which is positively correlated with fibrotic tissue formation and a lower level of apoptosis related to inflammation and cytotoxicity (Fig. 4p). Thus, it can be concluded that our new AoF-MSEA demonstrated excellent bio-compatibility and great potential in long-term implantable monitoring of intramuscular signals, owing to its tiny surgery incision, softness and stretchability.

In addition, another big advantage of AoF-MSEA is the multi channels electrodes distributed along the fiber, because one action always requires engagement of multi muscles and it is essential for the EMG prosthetic control to simultaneously monitor EMG signals from multiple muscles. These along-fiber channels can benefit the monitoring of multi muscles or motor units in muscles by only implanting one single fiber. But it is hard for rigid stainless-steel electrodes to construct along-fiber multi channels of electrodes and implanting an electrode in each muscle will cause severe tissue damage and an increased risk of infection. To demonstrate this, AoF-MSEA with two channels was sutured into the tibialis anterior and gastrocnemius muscles simultaneously, to monitor the rat’s hind leg movement. Distinct intramuscular EMG signals were recorded from the two muscles, affirming the fiber sensor’s capability to simultaneously obtain EMG information from different muscles (Fig. 4q, Fig. S38). Besides, as AoF-MSEA stretched with the muscle during the rat’s movement, it successfully captured mechanical strain via the sensor (Fig. 4r). Notably, since there would be slippage between AoF-MSEA and muscle especially at the beginning of the implantation, the deformation detected here should be smaller than that of muscle’s actual deformation (Fig. 4s). To further optimize this mechanical monitoring, the adhesive layer could be modified onto the surface of AoF-MSEA to enhance the synchronous deformation. Therefore, AoF-MSEA can achieve the long-term implantable monitoring of bio-electric signals and show great potential in bio-mechanical detection *in-vivo*.

## Conclusion

In summary, we report a new strategy to reduce the dimension of bioelectronics from 2D film to 1D fiber via a simple rolling method. By the topological geometry design, we fabricate more than 60-channel electrode array along one single microfiber and capacitive stretchable strain sensors in the same fiber. It is proved that 1D fiber AoF-MSEA significantly benefits the long-term implantable monitoring of bio-electrical and -mechanical signals owing to its tiny surgery incision, softness, and stretchability. It can continuously monitor the intramuscular EMG in rats for at least 43 weeks with more abundant frequency spectrum in the first 24 weeks. Also, it can act as a surgical suture into multiple muscles with an implantation incision of no more than 200 μm. After 13 months implantation, it has negligible effect on the surrounding muscles, especially compared to the rigid fiber with the same diameter. We show the evidence that the soft and stretchable 1D fiber shaped bioelectronics has superior properties in terms of the performance in long-term implantable monitoring in peripheral neuromuscular system. Additionally, by integrating a magnetic head, we highlight the impressive abilities of the soft microfiber electrode array to “submerge” into and “explore” the brain with minor surgery incision on skull. The proposed strategy provides a platform for the preparation and minimally invasive implantation of multifunctional fiber sensors. The proposed strategy not only opens up the study of multi-channel and -function soft and stretchable fiber-shaped sensors, particularly those with longitudinally distributed function units, but also establishes a minimally invasive implantation platform for biomedical and healthcare applications that require prolonged monitoring of physiological signals.

## Acknowledgments

we thank the Institutional Animal Care and Use Committee (IACUC) of Shenzhen Institute of Advanced Technology Chinese Academy of Sciences. All the animal experiments performed in this research were approved by the IACUC under the protocols of SIAT-IACUC-210316-JCS-XRJ-A1740-01.

## Funding

Shenzhen Science and Technology Program (KQTD20210811090217009)

National Natural Science Foundation of China (62101545 and 62201558)

National Key R&D Program of China (2021YFF0501601, 2021YFF0501602)

National Natural Science Foundation of China (62101544 and 62201559)

Guangdong Basic and Applied Basic Research Foundation (2020A1515110205)

The Fundamental Research Funds for the Central Universities (20720230041)

Partially supported by Shenzhen-Hong Kong Institute of Brain Science (NSY889021054) and the Science and Technology Program of Guangdong Province(2022A0505090007).

## Author contributions

R. Xie conceived of this research, performed the experiments, and wrote the manuscript. R. Xie, Q. Yu and F. Han performed the animal experiments and the ex vivo and in vivo EMG measurements, and J. Huang helped. F. Han and D. Li performed the implantation of the fiber sensor under magnetic force and the EEG measurements. X. Han and X. Xu helped to draw illustrations. X. Zhou provided the test equipment. Q. Tian, Q. Li, H. Li and W. Yan participated in the discussions. G. Li supervised the project. T. Xu and Z. Liu supervised the project and revised the manuscript.

## Data and materials availability

All data are available in the main text or the supplementary materials.”

## Supplementary Materials

Materials and Methods

Figs. S1 to S38

Movies S1 to S4

The code for processing electromyography (EMG) signals

