## Supplemental materials for "Rolling 2D Bioelectronic film into 1D: a Suturable Long-term Implantable Soft Microfiber"

Science
AAAS

### Supplementary Materials for

#### Rolling 2D Bioelectronic film into 1D: a Suturable Long-term Implantable Soft Microfiber

Ruijie Xie[1,2]†, Qianhengyuan Yu[1]†, Dong Li[3]†, Xu Han[2], Xiaolong Xu[2], Jianping Huang[1], Wei Yan[4]*, Xiaomeng Zhou[1], Xinping Deng, Qiong Tian[1], Qingsong Li[1], Hanfei Li[1], Hairong Zheng,[1] Guanglin Li[1], Fei Han[1]*, Tiantian Xu[3]*, Zhiyuan Liu[1]*

**The PDF file includes:**

Materials and Methods
Figs. S1 to S38
The code for processing electromyography (EMG) signals

**Other Supplementary Materials for this manuscript include the following:**

Movies S1 to S4

#### Materials and Methods

##### Fabrication of ultra-thin 2D film-shaped stretchable electrodes

Sodium polystyrene sulfonate solution (10 wt%) was spin-coated on a silicon wafer at 2000 rpm for 50 s to form a sacrificial layer. Polystyrene-block-poly(ethylene-ran-butylene)-block-polystyrene (SEBS) in toluene (7 wt%) was then spin-coated onto the PSS at 1000 rpm for 50 s to form a 400 nm thick layer of SEBS. The desired conductive patterns on the SEBS film were formed by thermal evaporation of gold under the cover of the mask. The evaporation rate was 0.5 Å/s. The thickness of the gold can be controlled by controlling the evaporation time.

##### Preparation of AoF-MSEA

A PET frame was first attached to the SEBS electrode. Then the silicon wafer was rinsed in water, the SEBS-based ultra-thin electrode was released from the silicon wafer after the sacrificial layer was dissolved by water. The ultra-thin electrode supported by the PET frame was picked up from the water and the gold film side was attached to the polytetrafluoroethylene (PTFE) board. The ultra-thin electrode was then rolled up carefully to form a microfiber sensor.

##### Integration of AoF-MSEA into an implantable system

The microfiber electrode was connected to signal acquisition equipment via a printed circuit board (PCB). The stainless-steel wire was welded to the PCB to connect it to the cranial interface. The anisotropic conductive film was then attached to the PCB. Next, the wire sensor was attached to the anisotropic conductive films to connect it to the PCB, and the PCB was wrapped with a parafilm to protect the interface from subsequent operations. Finally, the PCB was sealed with DOWSIL™ 734 Flowable Sealant.

##### Minimally-invasive implantation of AoF-MSEA

Three-month-old SD rat was used to assess the performance of long-term monitoring of AoF-MSEA. A 1 cm wound was made in the rat's waist and the PCB was fixed in the wound of the waist to prevent it from moving. Then one end of the surgical threads was attached to the puncture needle and the other end to AoF-MSEA. After the needle was passed through the muscle from the wound in the waist to the leg, the wire sensor was brought in for minimally invasive implantation and the surgical thread was fixed in the muscle. A 1cm diameter wound was then made in the skull, the connector was passed subcutaneously from the waist wound to the head wound and fixed in the skull with screws. Finally, the wound on the skull was closed with dental cement and the wound in the waist was sutured with surgical threads. Intramuscular EMG signals and strain signals were recorded from the skull connector. For long-term EMG monitoring, the Intramuscular EMG signal was recorded every two weeks during the rat's recovery period after surgery. After three weeks, the Intramuscular EMG signal was recorded once a week until the device failed. Tissue staining was performed by Shanghai Boerfu Biotechnology Co. Ltd. This experiment was approved by the Institutional Laboratory Animal Management and Use Committee of Shenzhen Institute of Advanced Technology, Chinese Academy of Sciences. The project number was SIAT-IACUC-210316-JCS-XRJ-A1740-01.

In detail, from August 16, 2021, to January 6, 2022, we conducted a series of long-term implantation experiments on five rats (designated as #201, #205, #209, #217 and #220), involving AoF-MSEA, 2D thin-film electrodes, and stainless steel wire electrodes. The primary implantation sites were the anterior tibialis and gastrocnemius muscles on both sides of the rats. Detailed information about the implanted rats, including their identification numbers, implantation dates, implantation locations, electrode types, and more, is provided in Table S1.

Among these rats, four completed data collection for over three months, while one rat (#205) died 59 days after the implantation.

The key experimental process began with the anesthesia of the rats using isoflurane anesthesia. Subsequently, the fur was shaved from the lateral side of the hind thigh, extending to the femur region, as well as from the back and skull areas. An injection needle was then used, following the direction of muscle fibers, to insert the fiber electrode along the tip of the needle into the needle cavity, which was gradually withdrawn from the muscle bundle. This left the fiber electrode embedded within the muscle. To secure the electrode, the end of the fiber electrode was sutured to the muscle fascia using surgical thread. The soft-hard interface was extended subcutaneously along the rat's back and fixed to the grounding part of the electrode. The rear end of the soft-hard interface was extended subcutaneously to the skull connection point via a stainless steel wire. After disinfecting the skull surface with an alcohol swab, the skin over a 1.5 cm × 1.0 cm area of the brain was removed, and the skin tissue was cleaned in the front to fully expose the skull. The skull connection point, with the mother head facing upward, was placed in the middle of the fixture. Using a flat-headed 1mm × 3mm screw, the fixture's position was secured on the skull surface through the fixture's holes (position 14 in Fig. S18). A relatively diluted dental cement mixture, consisting of dental cement and dental cement powder, was introduced through the fixture's holes (position 21 in Fig. S18) to fill the dental cement around the fixture, covering the entire fixture and part of the skull interface. The dental cement was allowed to cure, thus completing the preparation of the skull connection point. The primary procedure for fiber electrode implantation is as described. For stainless steel wire electrodes, the surgical process was essentially the same, involving the fixation of the stainless steel wire electrode to the muscle fascia using sutures. In the case of film electrodes, there was a larger incision area, requiring multiple fixation points.

**Bio-signals collection procedures during Long-term monitoring of AoF-MSEA**

After electrodes implantation, impedance data at rest and electromyographic data during running were collected weekly for each rat. Some data collection intervals were affected due to the pandemic (COVID-19). Prior to post-implantation dissection, X-ray imaging was performed to assess the condition and morphology of the electrodes.

#201 rat (implanted on August 16, 2021, died on September 24, 2022, and sampled) had a stainless-steel wire fracture at the cranial connector joint, and the fiber electrode broke into two parts. Visible to the naked eye, there was a soft/hard interface water ingress. Using exclusionary methods and testing channel-to-channel impedance, lower values indicated poor sealing at the soft/hard interface for this rat.

#209 rat (implanted on October 23, 2021, died on April 23, 2022) had a membrane electrode that had shifted from its original implantation site, with instability observed at the soft/hard interface.

#220 rat (implanted on January 6, 2022, died on September 25, 2022, and sampled) had a stainless-steel wire fracture at the cranial connector joint. However, the electrode and the soft/hard interface remained intact, and testing revealed good sealing at the soft/hard interface.

#217 rat (implanted on November 24, 2021, died on December 1, 2022, and sampled) had an intact soft/hard interface and cranial connector joint, making it of significant research value. Consequently, we conducted a detailed 10-month analysis of the data from this rat, which includes impedance (Fig. 4h), electromyographic data (Fig. S22), and staining analysis (Fig. 4i-4p).

**EEG Collection in rats**

The subject used was male Sprague-Dawley (SD) rats weighing around 900 grams. A hole with a diameter of 1mm was drilled above the rat's skull using a drilling device. With the assistance of ultra-fine stainless-steel needles (with a diameter of smaller than 10 micrometers), AoF-MSEA was carefully inserted through the drilled hole into the rat's brain under the magnetic field. Due to the relatively small size of the rat's brain, this experiment simultaneously collected brainwave signals from two channels of the rat.

**Demonstration of AoF-MSEA wandering in an artificial brain under magnetic field**

A model of the cerebral vasculature was made using 3D printing technology, and an agar solution with a mass fraction of 0.5% was poured into this model. After the agar cured at room temperature, an agar-filled cerebral vasculature model was obtained. A stretchable wire sensor with a magnetic needle fixed at one end was placed in the model, and the external magnet was used to attract the needle, which drove the movement of AoF-MSEA in the cerebral vascular model. The position and direction of the external magnet was repeatedly adjusted to guide and advance the stretchable wire electrode to avoid the blood vessel. In addition, five agar gels with different simulated signals were used to simulate neural units. According to the above method, a stretchable wire sensor with a magnetic needle was driven by an external magnet and moved along a linear path or a curved path. The signal applied to the gel can be detected as the wire passes through the gel.

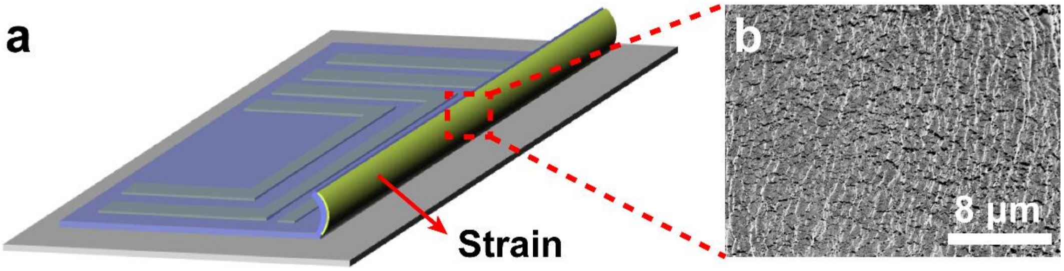


**Fig. S1. Morphological changes of Au film during rolling process.** a) Illustration of the film electrode being rolled up from the substrate. b) SEM image of the film electrode after being rolled up.

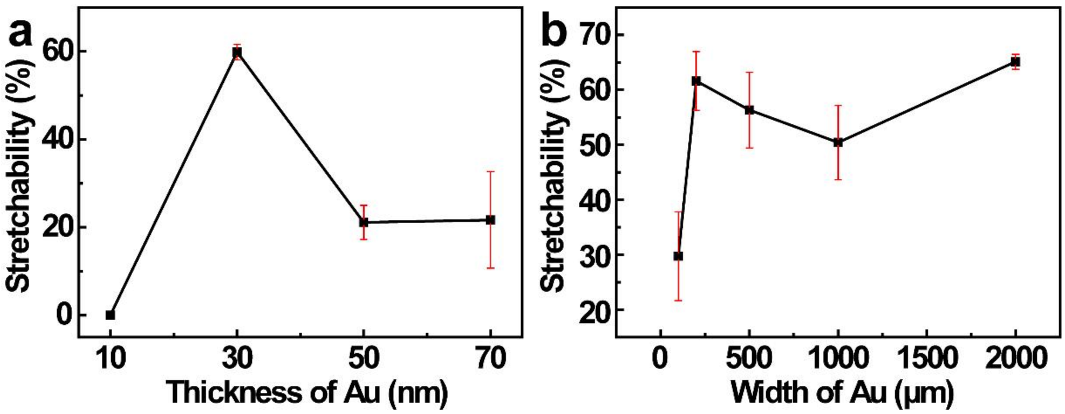

**Fig. S2. Stretchability optimization of thin film electrodes.** a) For a given width of conductive path, the stretchability of film electrodes with different thicknesses of Au. b) For an Au thickness of 30 nm, the stretchability of film electrodes with different of widths of conductive path.

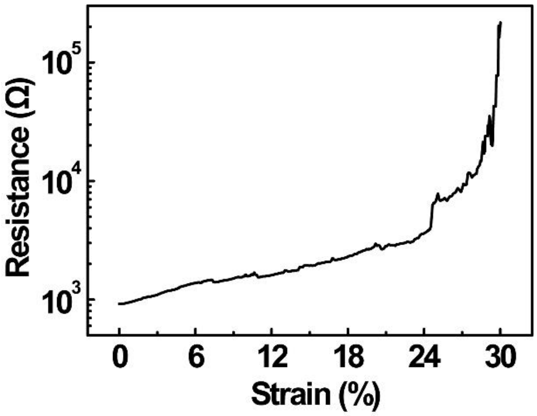

**Fig. S3.** The stretchability of the 60-channel fiber electrode.

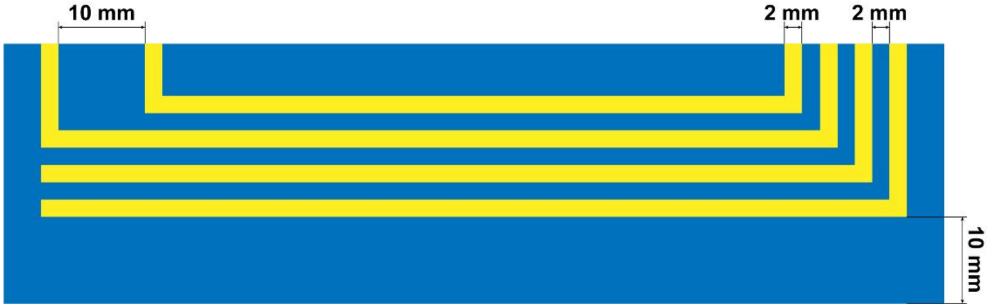


**Fig. S4.** The patterns design of the film electrode.

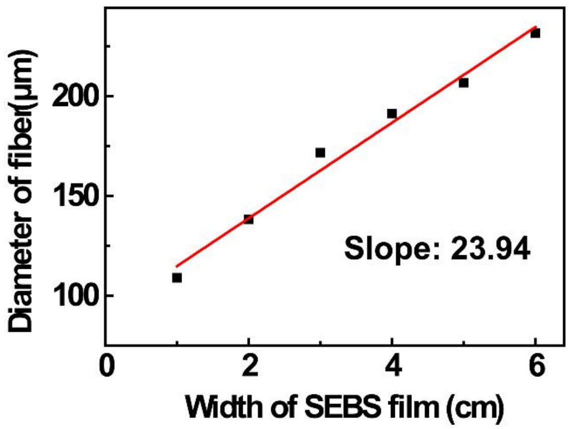

**Fig. S5.** Diameter of fiber devices fabricated from thin film electrodes of different widths.

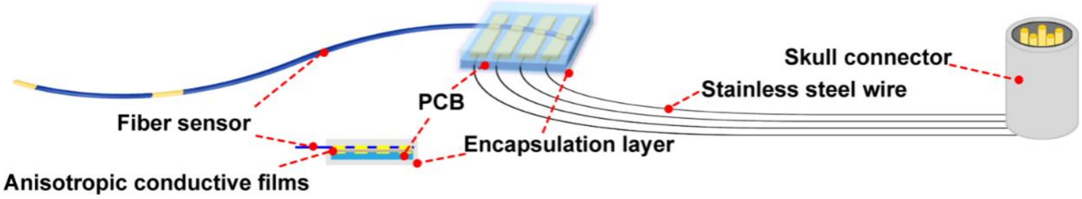

**Fig. S6.** Interface design of fiber sensor to signal acquisition devices.

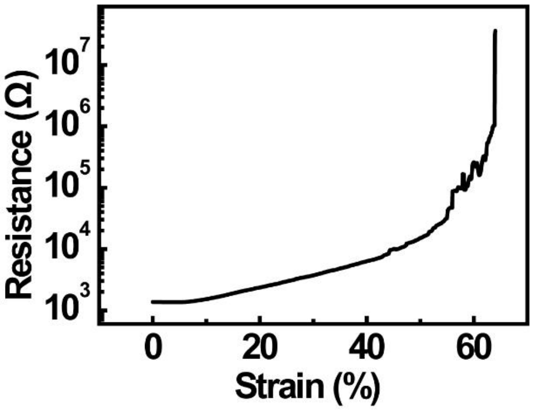

**Fig. S7**. The first tensile test of the four-channel fiber electrode upon fabrication.

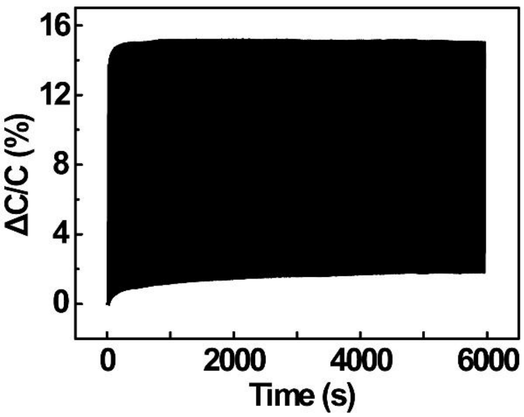


**Fig. S8.** The performance of the fiber capacitive strain sensor after being stretched to 20% for 1000 cycles.

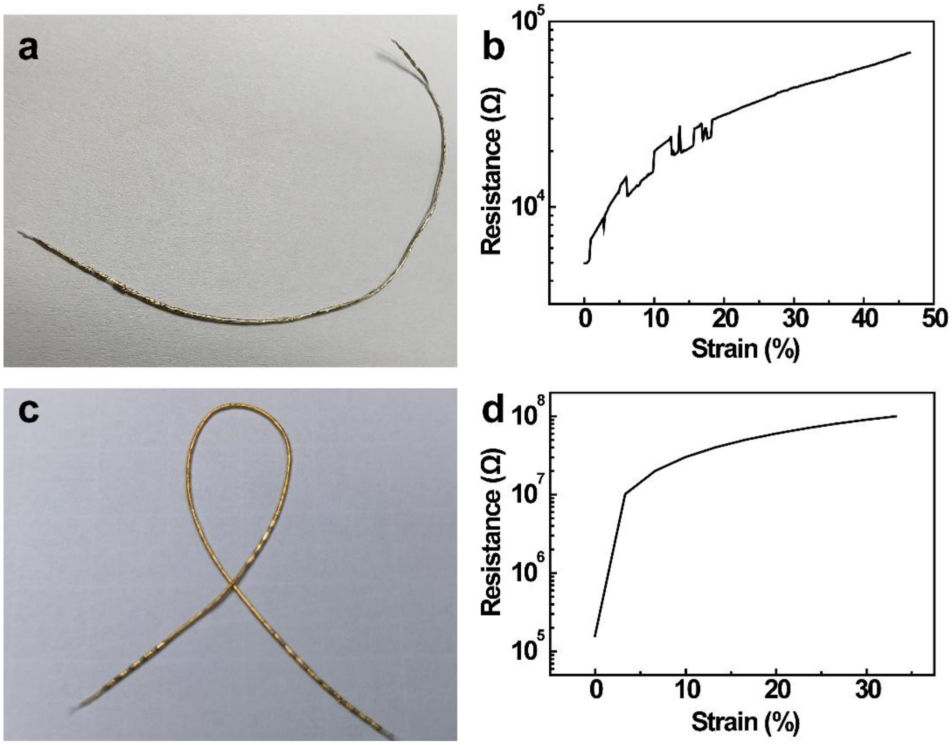


**Fig. S9. Universality of the rolling strategy.** a) Photo of the ethylene-vinyl acetate copolymer (EVA)-based fiber sensor. b) Stretchability of the EVA-based fiber sensor system. c) Photo of the polydimethylsiloxane (PDMS)-based fiber sensor. d) Stretchability of the PDMS-based fiber sensor system.

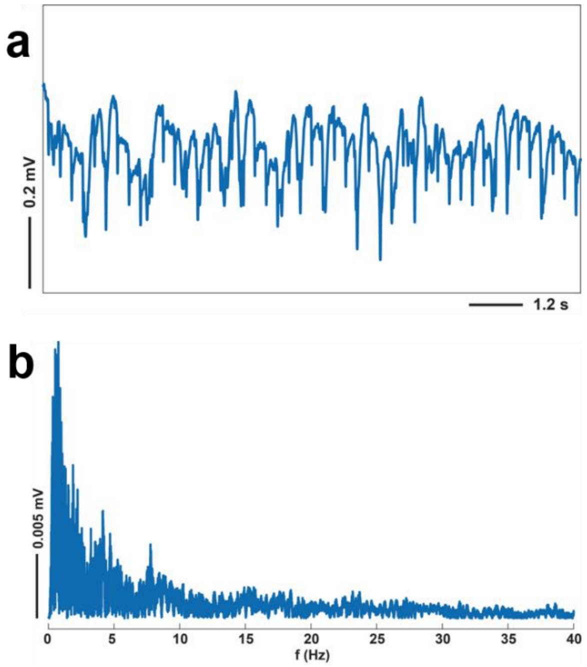


**Fig. S10. EEG monitoring.** a) Electrical signals recorded by fiber electrodes in anesthetized rats. b) Spectrum diagram of the electroencephalogram.

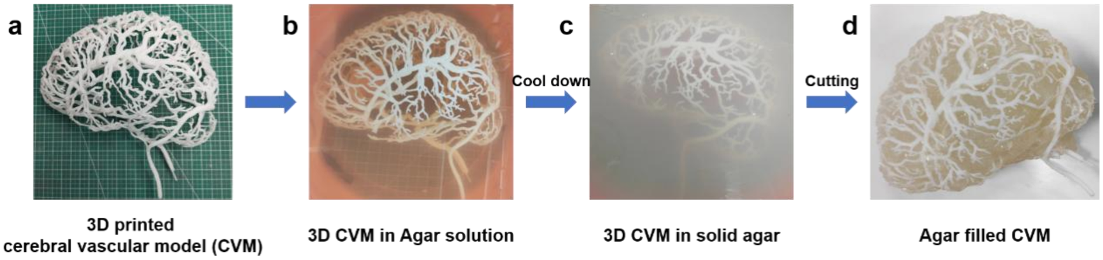


**Fig. S11. Simulation of human cerebral vascular model.** (a-d) The flow diagram of three dimensional (3D) printed cerebral vascular model filled with 0.5% wt agar.

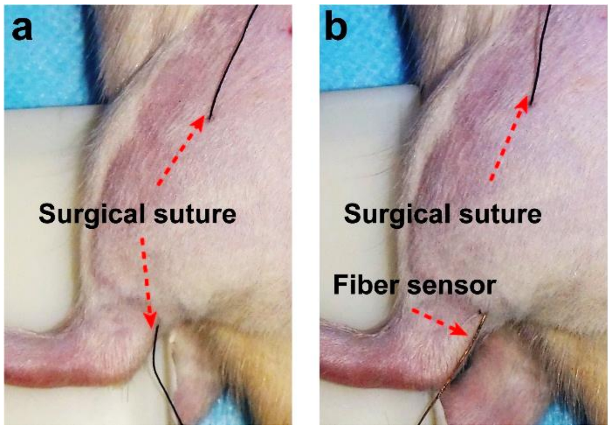

**Fig. S12.** a-b) The process of sewing the wire sensor into the muscle.

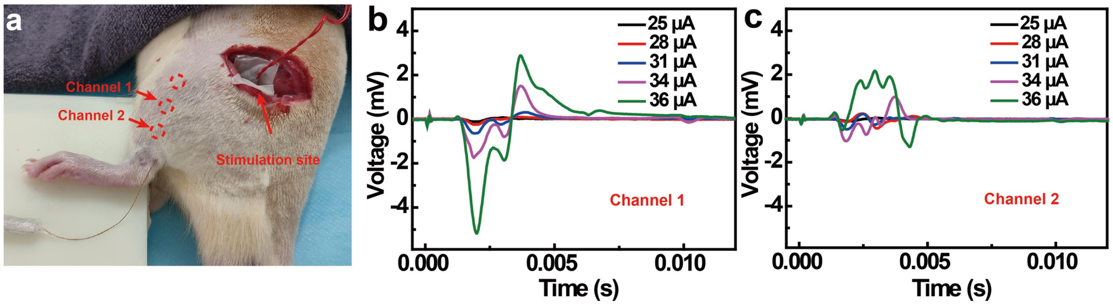


**Fig. S13. Measurement of intramuscular EMG induced by stimulation through a wire sensor.** a) Wire sensor monitoring myoelectricity evoked by sciatic nerve stimulation. b) EMG signals excited by different stimulation currents detected by channel 1. c) EMG signals excited by different stimulation currents detected by channel 2.

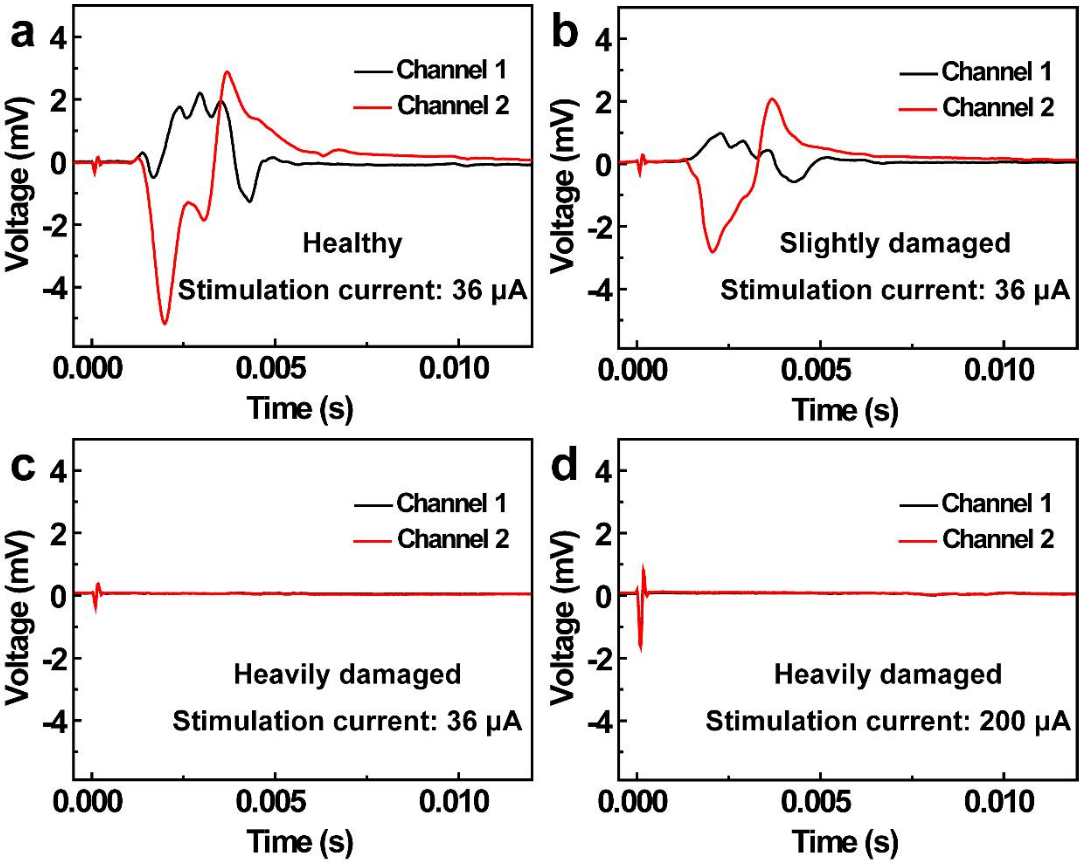


**Fig. S14. AoF-MSEA as a tool for assessment of the extent of nerve damage.** a-c) EMG signals evoked by stimulation of the sciatic nerve at different levels of sciatic nerve injury (a) healthy; (b) slightly damaged; (c) heavily damaged. d) EMG signals evoked by a stimulation current of 200 μA when the sciatic nerve was severely damaged.

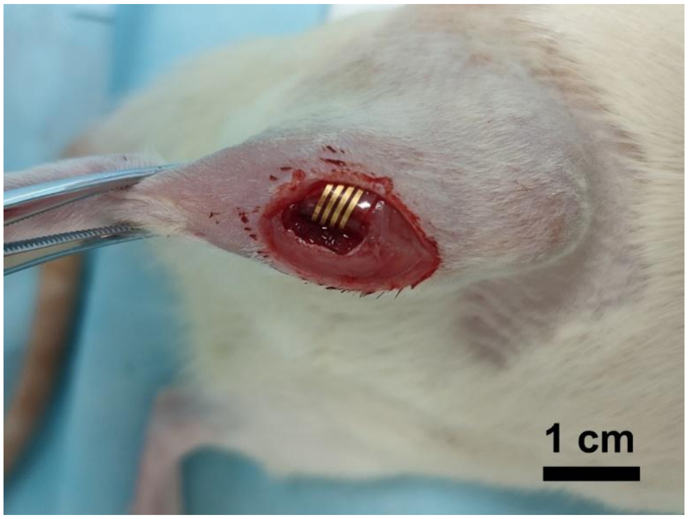


**Fig. S15.** The wound created for the implantation of the 4-channel film electrode.

**Table S1**. The detailed information on the individual implantation of electrodes in five rats.

| Rat ID | #201 | #205 | #209 | #217 | #220 |
|---|---|---|---|---|---|
| **Implantation plan** | AoF-MSEA in tibialis anterior muscle (TAM) | AoF-MSEA and stainless steel wire electrode in same TAM | Film electrode in TAM | AoF-MSEA in Gastrocnemius (GAST) and TAM; Stainless steel wire electrode in TAM | AoF-MSEA (ten channels) in GAST and TAM |
| **Implantation date** | 2021/8/16 | 2021/9/23 | 2021/10/23 | 2021/11/24 | 2022/1/6 |
| **Days Since Implantation** | 404 | 61 | 182 | 372 | 262 |
| **Longest EMG Day (Post-Implantation)** | 382 | 59 | 177 | 304 | 261 |
| **Anatomical results** | 1. Stainless steel wire breaks at skull interface 2. Electrode breakage 3. Tissue fluid infiltration in soft-hard interface | - | 1.Implant site displacement 2. Tissue fluid infiltration in soft-hard interface | 1 Intact skull interface and soft-hard interface 2. Electrode breakage | 1. Stainless steel wire breaks at skull interface 2. Intact soft-hard interface |

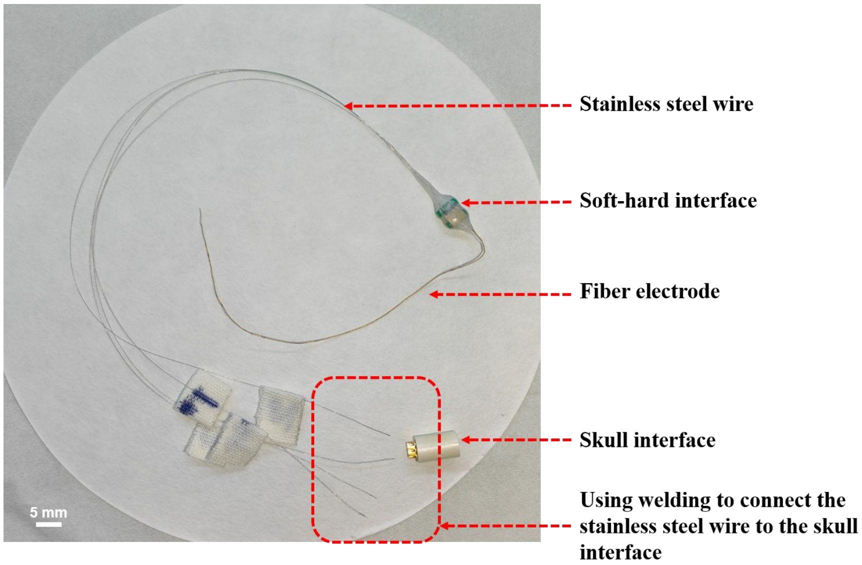


**Fig. S16.** Comprehensive structural diagram of the microfiber electrode system, encompassing the fiber electrode, the soft-hard interface, and the welding to the skull interface through stainless steel wire.

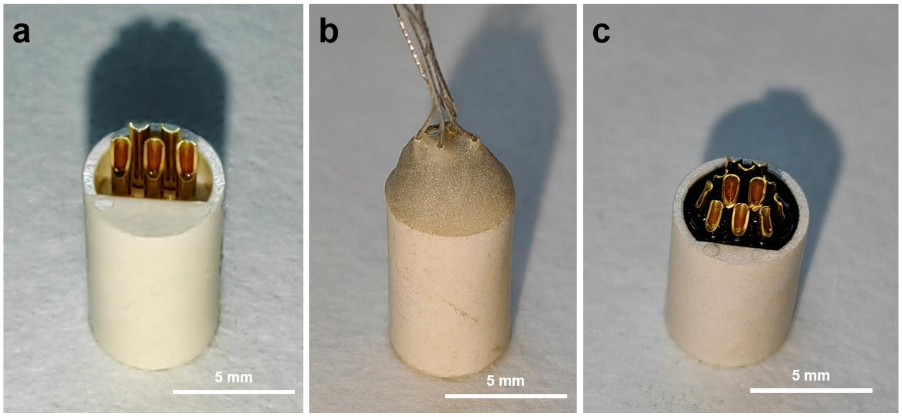


**Fig. S17.** a) Five-channel skull interface (unassembled components). b) Five-channel skull interface (assembled with connected wires and sealed using dental cement). c) 12-channel skull interface.

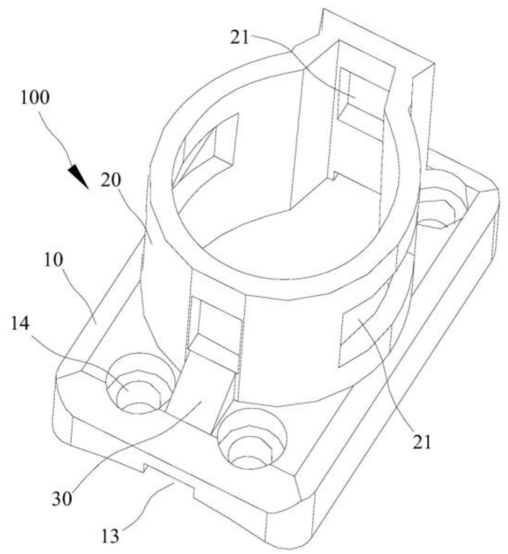


**Fig. S18.** Illustration of the skull interface fixture.

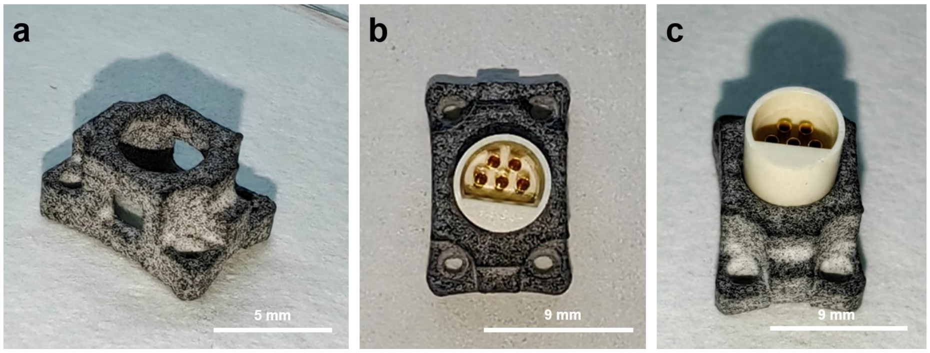

**Fig. 19.** a) Photo of the skull interface fixture. b-c) Top view and side view of the schematic diagram of the skull interface in the fixture.

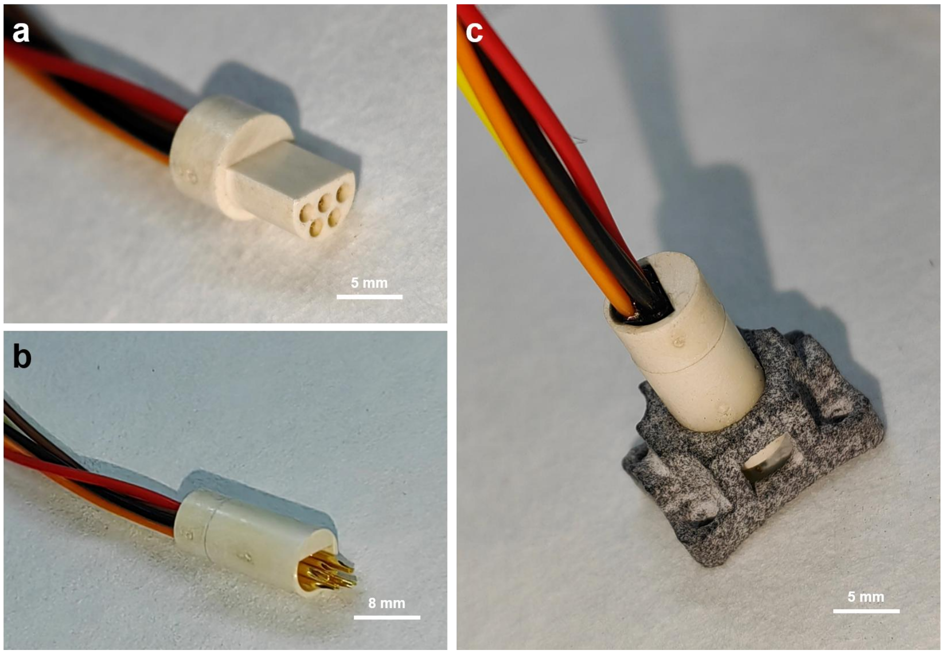

**Fig. 20.** Schematic diagram of the connection between the male and female parts of the skull interface used for data collection.

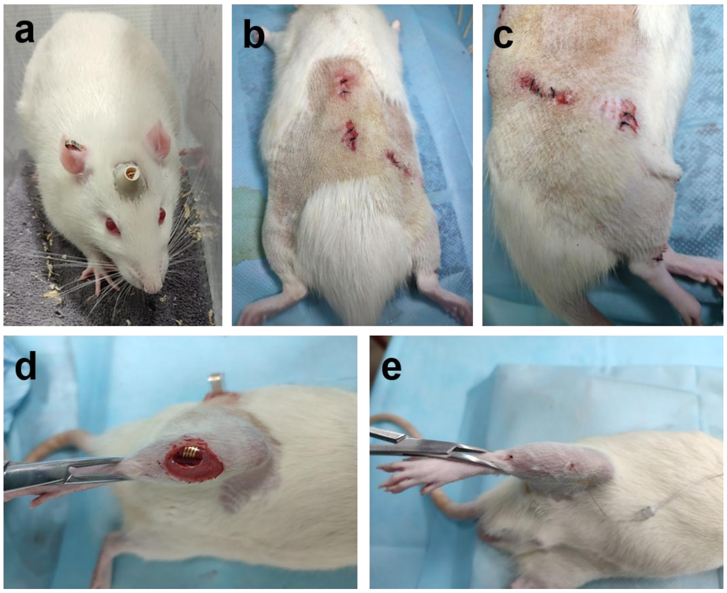


**Fig. 21.** a) Rats after completion of the skull interface surgery. b-c) Sutures on the wound after implanting the fiber electrode in the rat. d) Illustration of thin-film electrode implantation. e) Illustration of the minimally invasive implantation of the fiber electrode into the muscle.

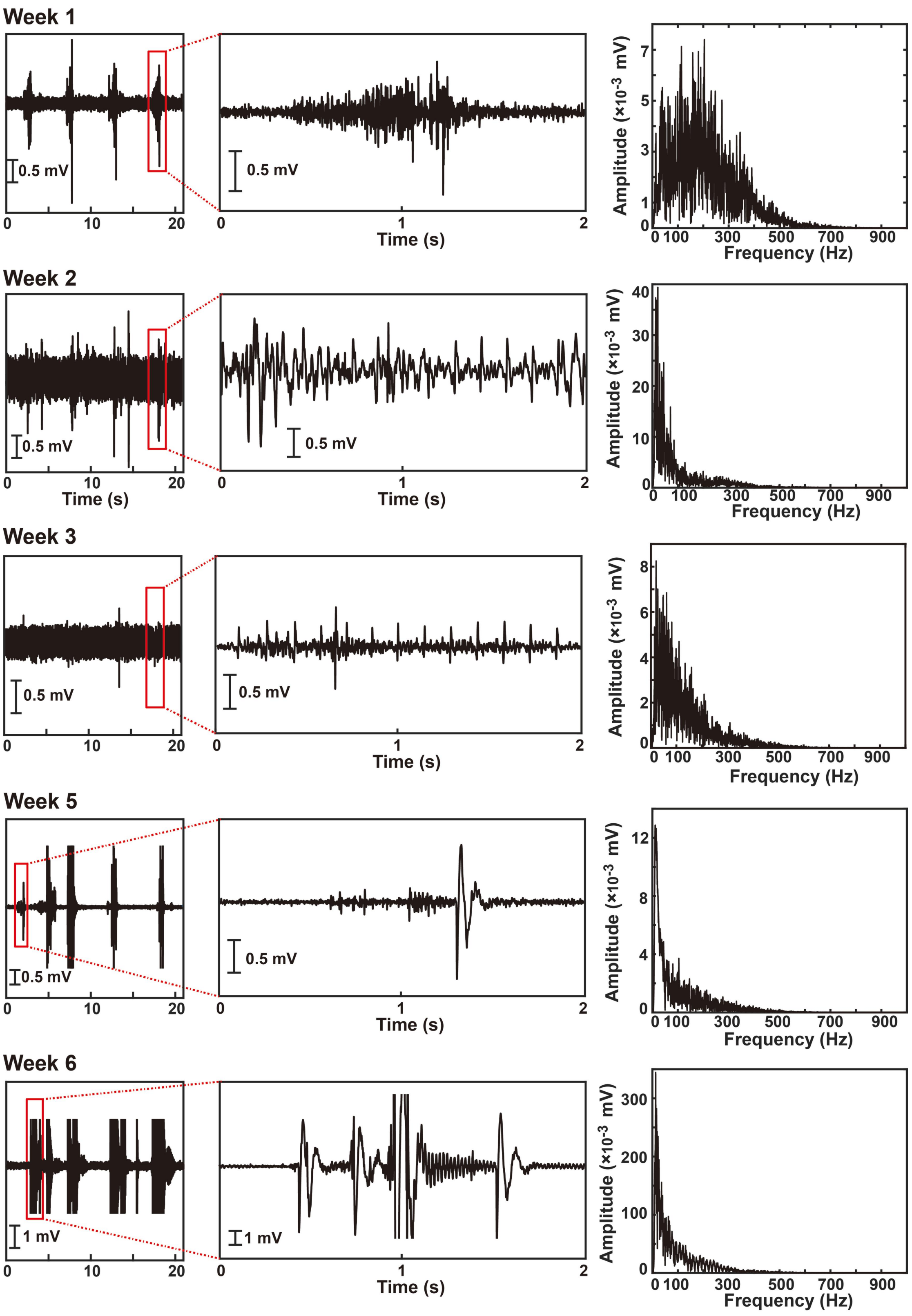
Week 1
0.5 mV
0
10
20
0.5 mV
0
1
2
Time (s)
Amplitude (×10^-3 mV)
7
5
3
1
0
0 100
300
500
700
900
Frequency (Hz)
Week 2
0.5 mV
Time (s)
40
30
20
10
Week 3
8
6
4
2
Week 5
12
8
4
Week 6
1 mV
300
200
100

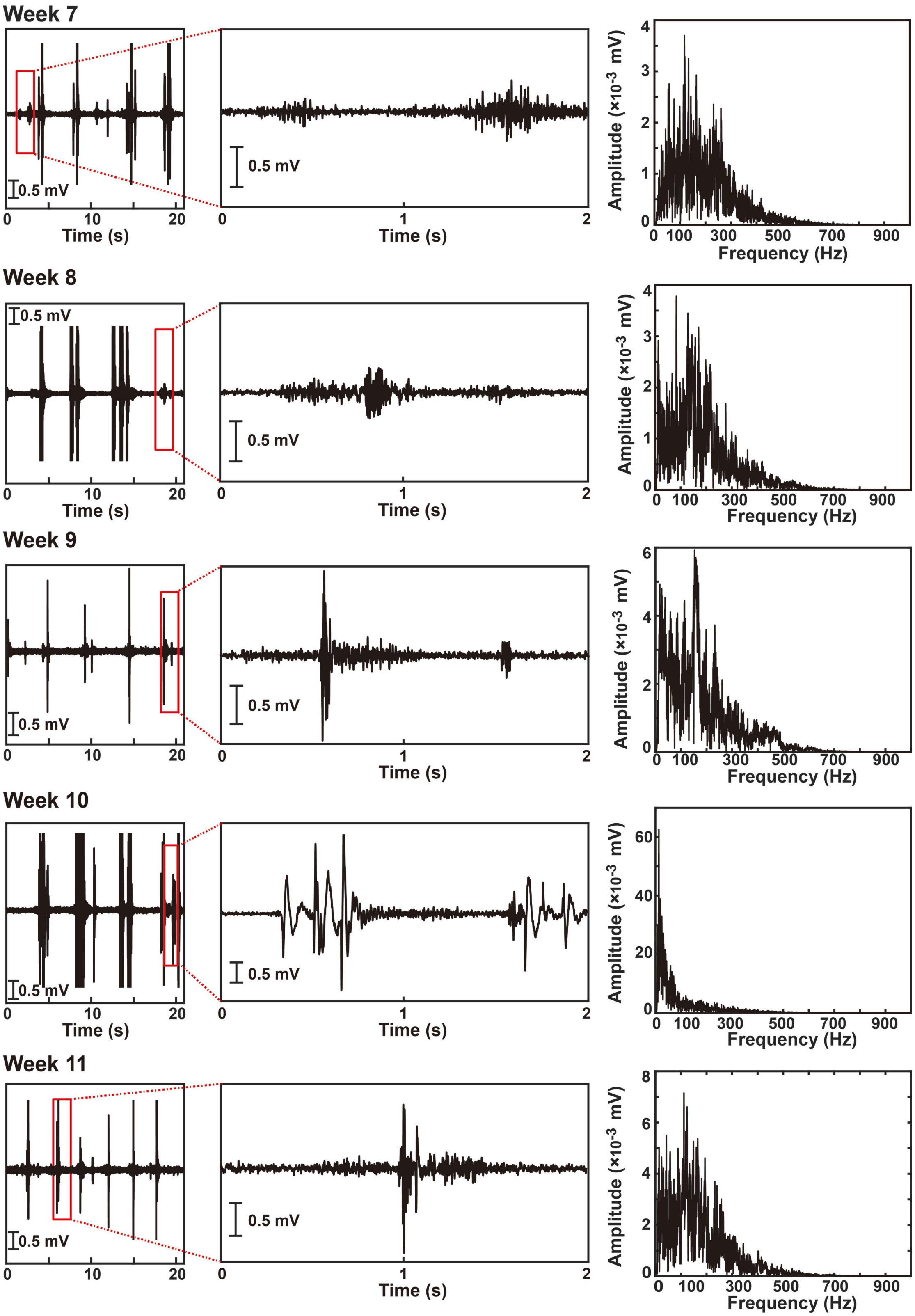

Week 7
0.5 mV
Time (s)
Amplitude (×10-3 mV)
Frequency (Hz)
Week 8
Week 9
Week 10
Week 11

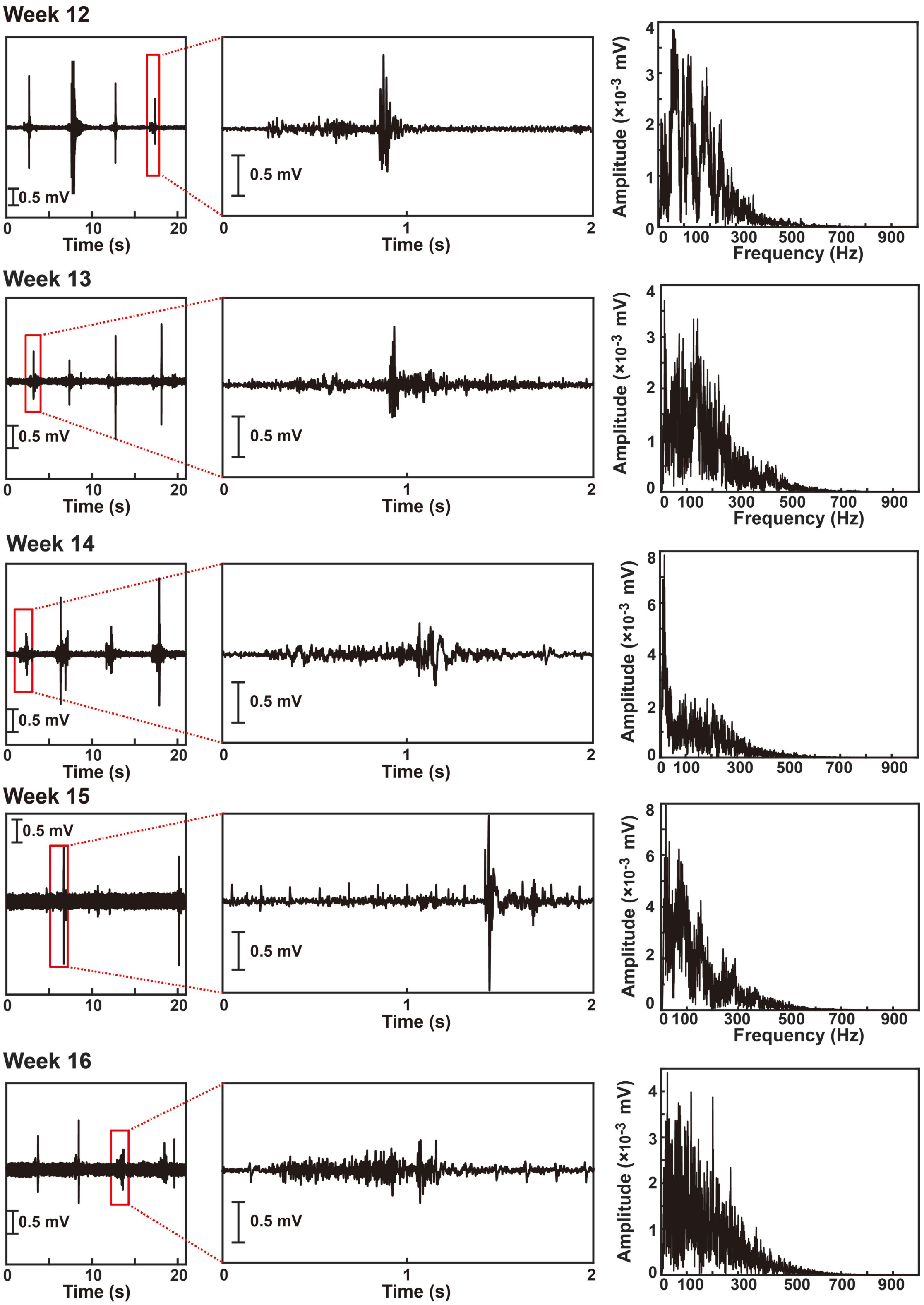
Week 12
0.5 mV
Time (s)
Amplitude (×10$^{-3}$ mV)
Frequency (Hz)
Week 13
0.5 mV
Time (s)
Amplitude (×10$^{-3}$ mV)
Frequency (Hz)
Week 14
0.5 mV
Time (s)
Amplitude (×10$^{-3}$ mV)
Week 15
0.5 mV
Time (s)
Amplitude (×10$^{-3}$ mV)
Frequency (Hz)
Week 16
0.5 mV
Time (s)
Amplitude (×10$^{-3}$ mV)

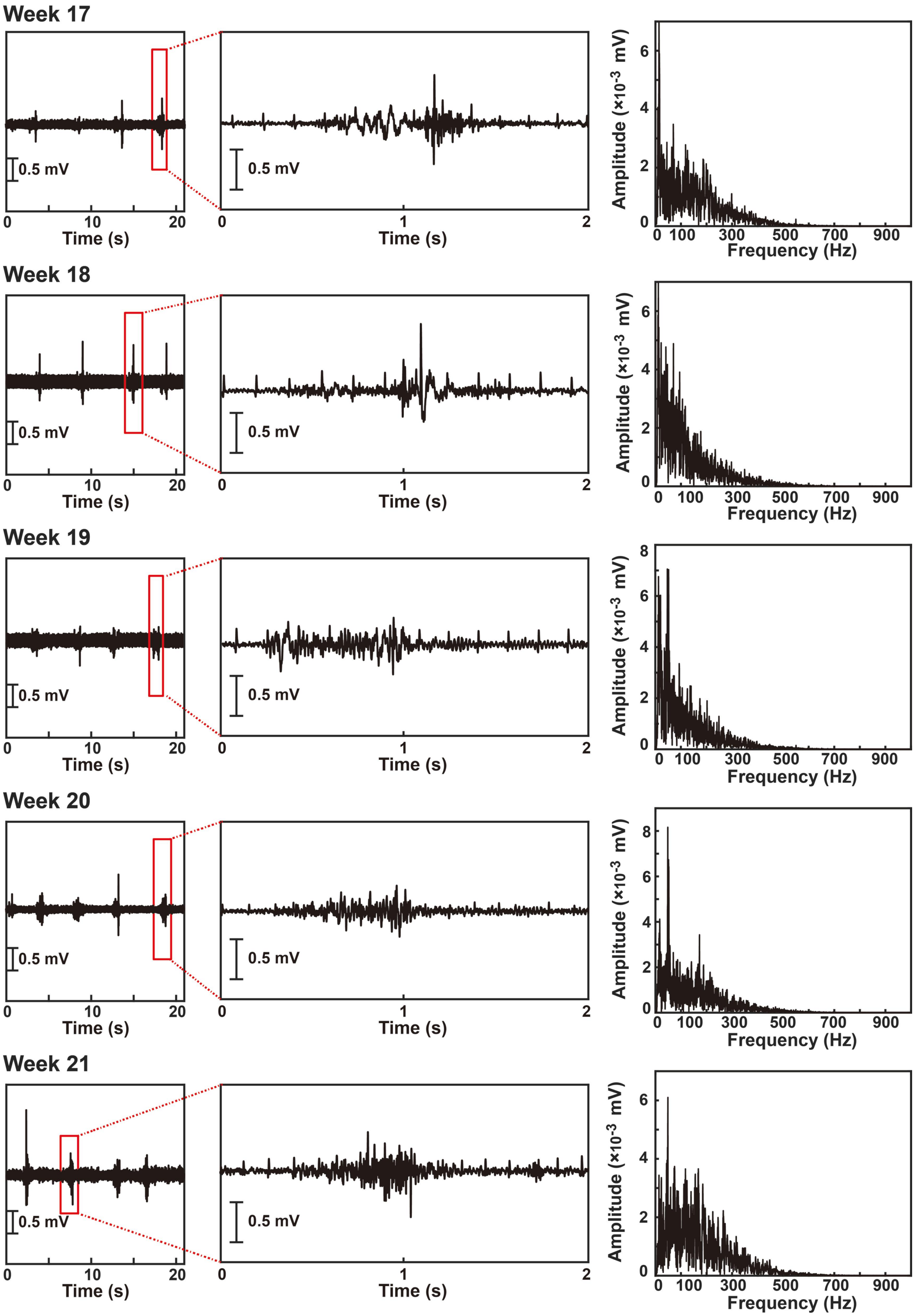
Week 17
Week 18
Week 19
Week 20
Week 21
0.5 mV
Time (s)
Amplitude (×10-3 mV)
Frequency (Hz)

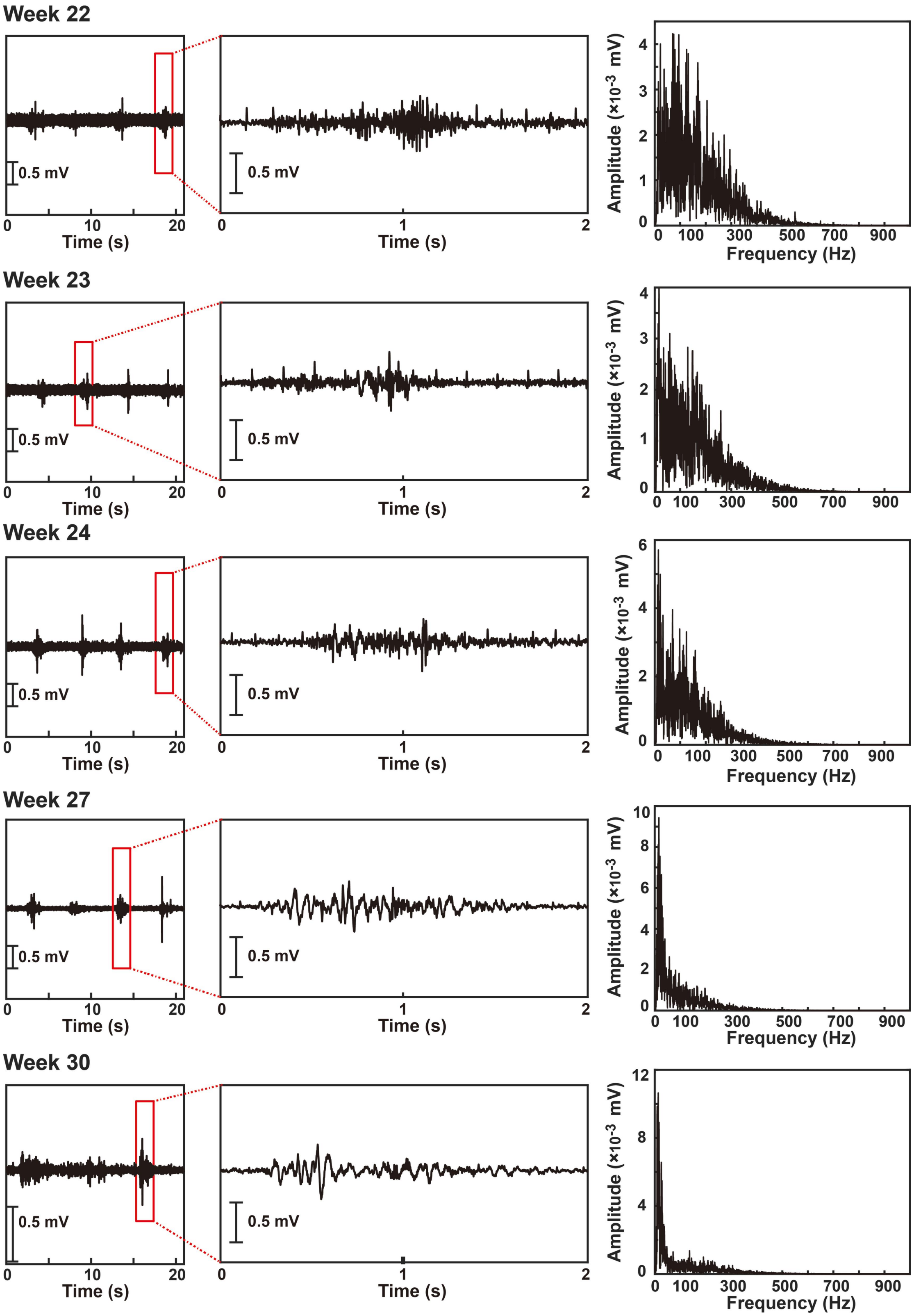
Week 22
0.5 mV
0.5 mV
Time (s)
Amplitude (×10$^{-3}$ mV)
Frequency (Hz)
Week 23
0.5 mV
0.5 mV
Time (s)
Amplitude (×10$^{-3}$ mV)
Week 24
0.5 mV
0.5 mV
Time (s)
Amplitude (×10$^{-3}$ mV)
Frequency (Hz)
Week 27
0.5 mV
0.5 mV
Time (s)
Amplitude (×10$^{-3}$ mV)
Frequency (Hz)
Week 30
0.5 mV
0.5 mV
Time (s)
Amplitude (×10$^{-3}$ mV)
Frequency (Hz)

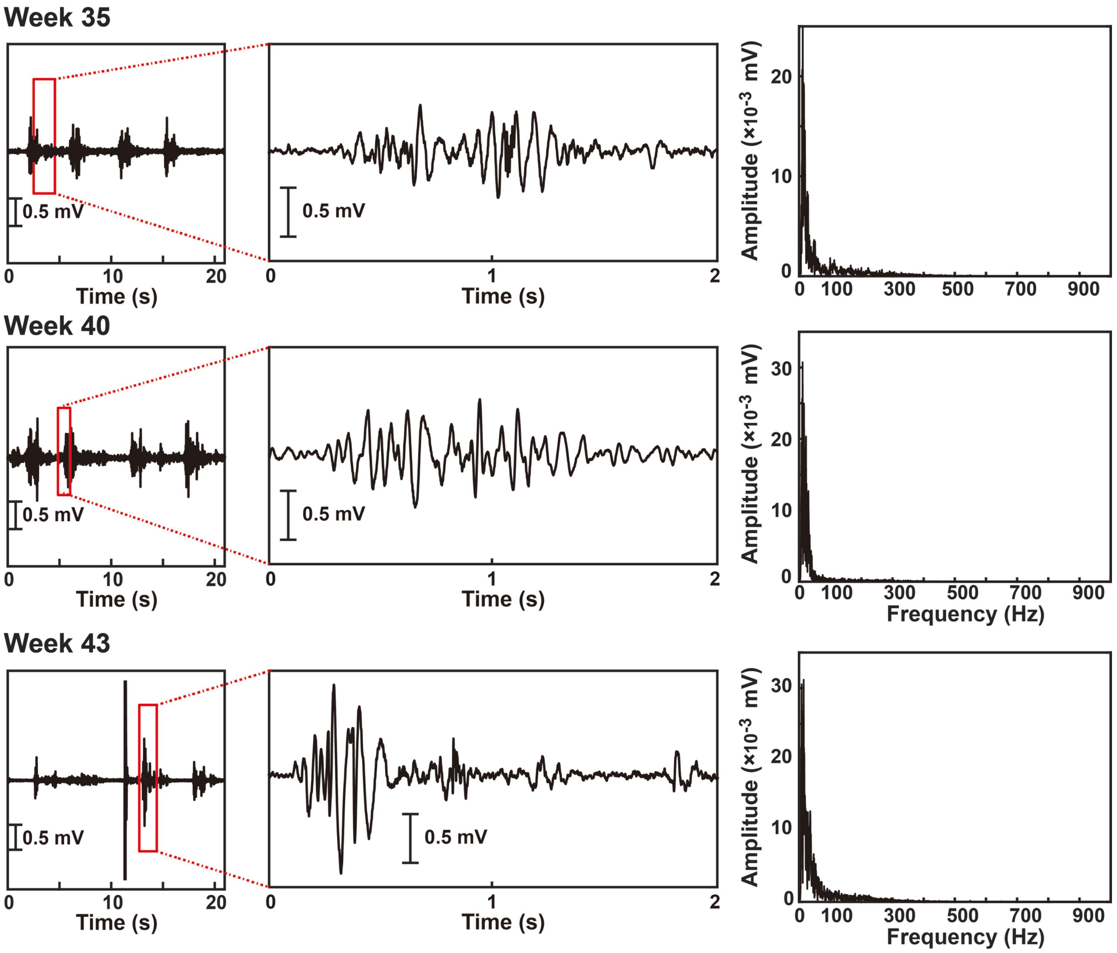

**Fig. S22.** 43-week EMG data for #217 rat. From left to right, the weekly data includes the EMG graph, the enlarged view, and the corresponding spectral graph for enlarged view.

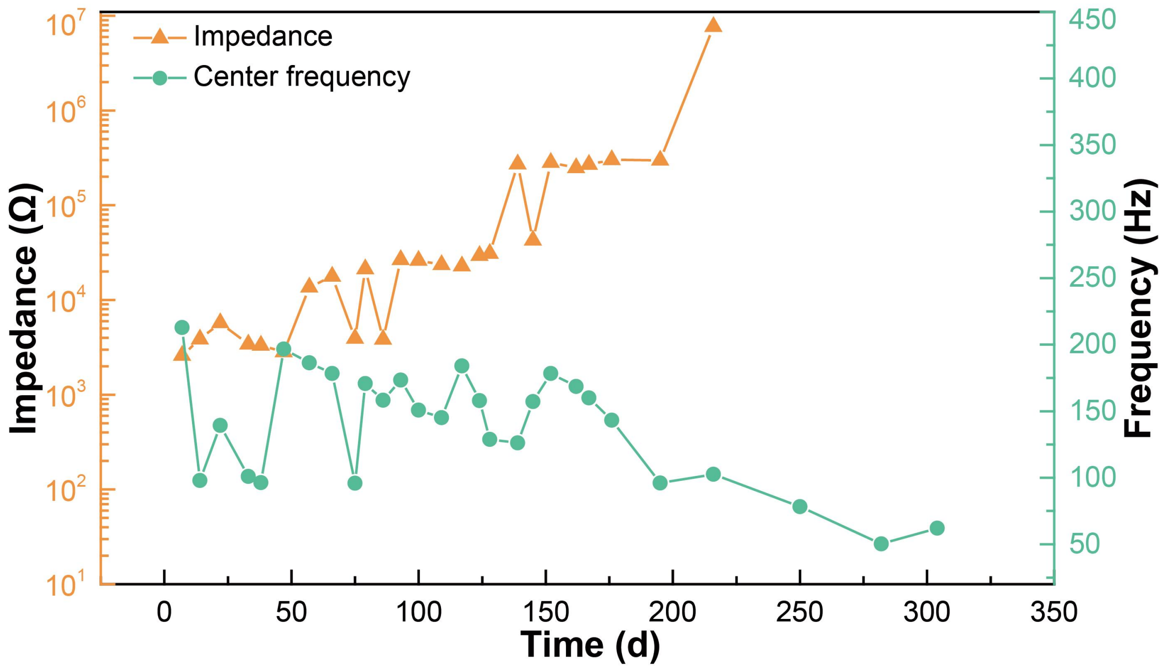


**Fig. S23.** Changes in electrode impedance and fluctuations in electromyographic central frequency values corresponding to the data at 43 weeks of implantation

**The major anatomy findings and conclusions for #201, #209, #217, and #220 are as follows:**

**#201 (13 months implantation, conducting dissection experiments in the 57th week.)**

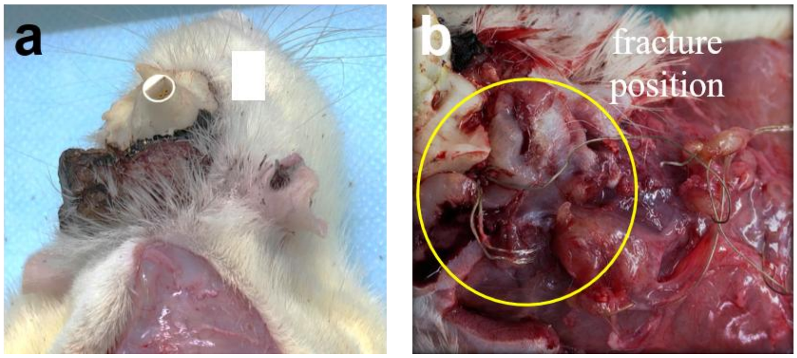


**Fig. S24.** a) During electrode implantation (June 7, 2022), there was partial displacement of the stainless steel wire at the skull interface. The rats were anesthetized, and the stainless steel wire at that location was adjusted. Suturing was performed to minimize the wound, and dental cement was used for fixation. b) During the dissection, tissue overgrowth was observed in the area, and upon examination, it was confirmed that the stainless steel wire had fractured, with only one intact connection remaining.

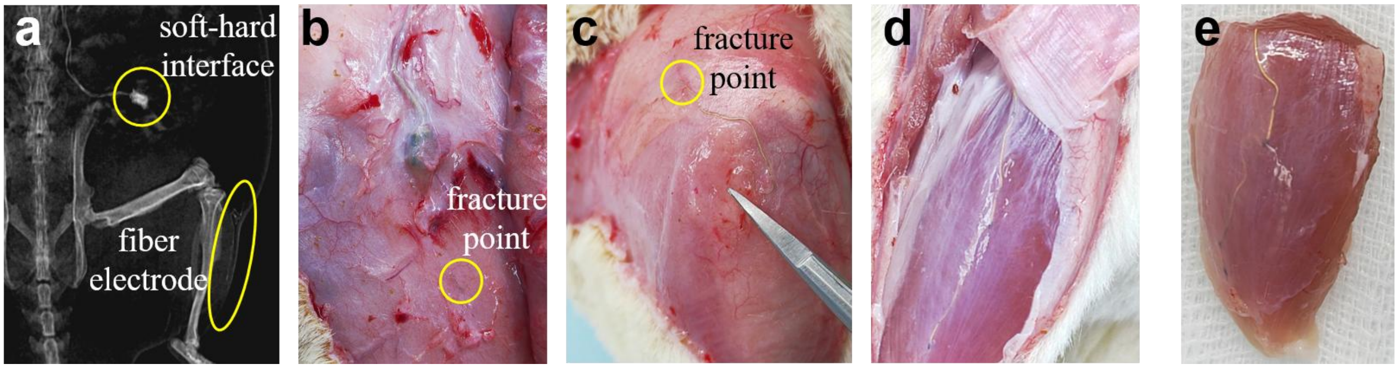


**Fig. S25.** a) X-ray observations revealed a breakage between the fiber electrode and the soft-hard interface. b-c) Anatomy images show the two ends of the electrode fracture point. d-e) The fiber remains intact within the muscle bundle.

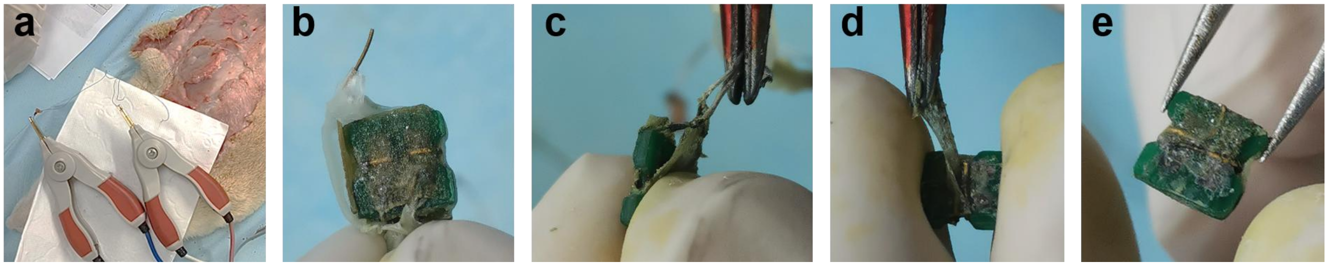


**Fig. S26.** a) Impedance schematic diagram between channels during *in-vivo* testing. b-e) Disassemble the soft-hard interface and visually inspect the situation for evidence of water corrosion, demonstrating its unstable sealing integrity.

**#209**

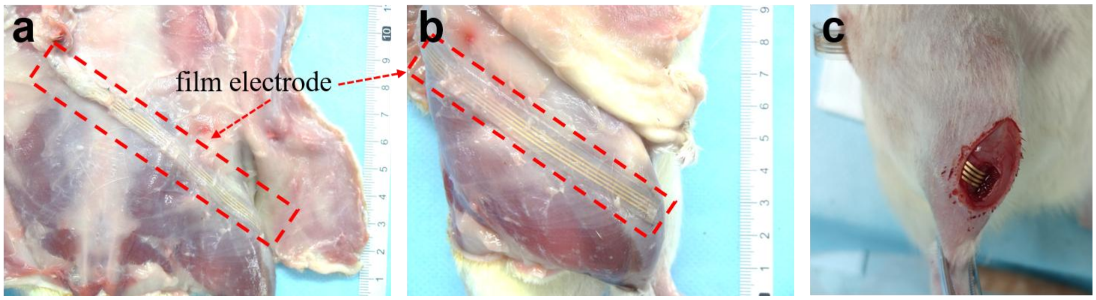


**Fig. S27.** a-b demonstrates the position of the film electrode during dissection, while c shows the position during implantation. It is evident that the film electrode has shifted compared to its implantation position, with the measuring point moving from within the muscle to outside the muscle fascia.

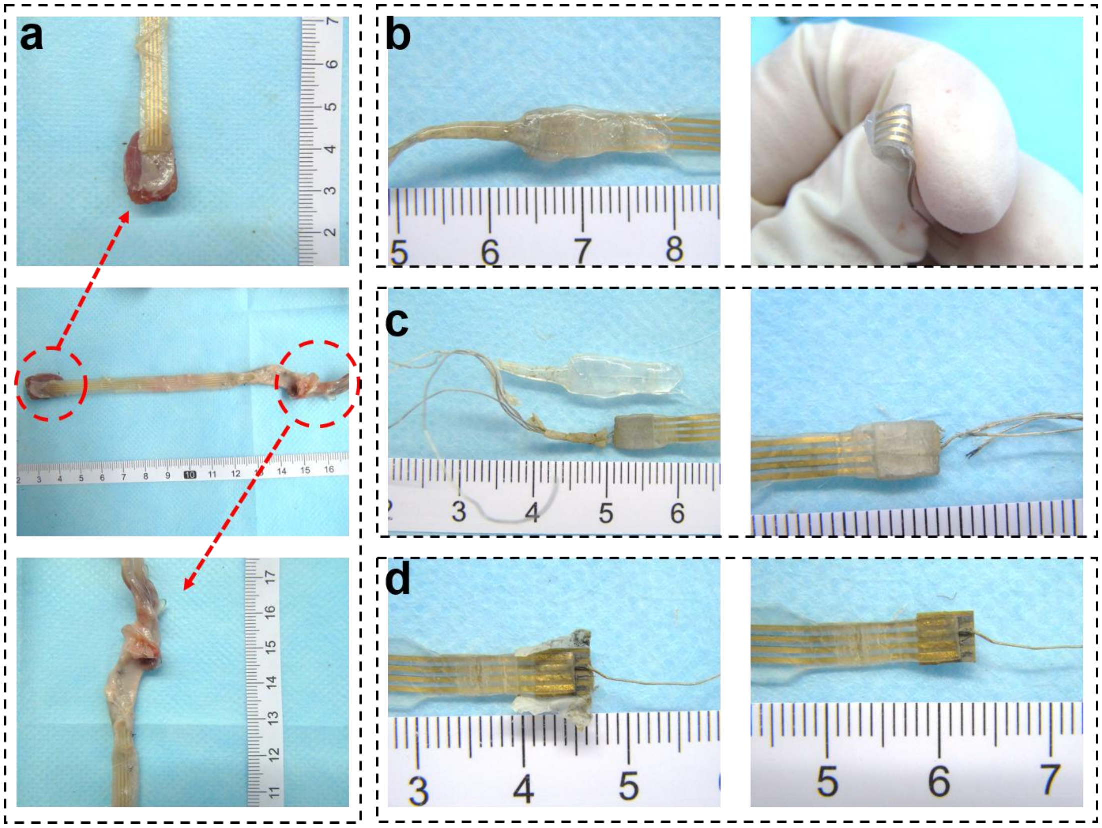


**Fig. S28.** a) During the sampling process, the entire electrode device was removed. The electrode had 4 channels for measurement, with two channels reserved for tissue fixation, and the remaining channels were used for optical observation of their surface morphology. Notably, there was visible tissue wrapping and accumulation around the stainless steel wires. b) There is a gap between the encapsulation layer of the soft-hard interface and the electrode layer. c-d) Disassembling the soft-hard interface revealed that the stainless steel wire leads used for welding had come loose. The likely cause of this issue is that tissue fluid entered the interface through the gap shown in Fig. b and led to corrosion.

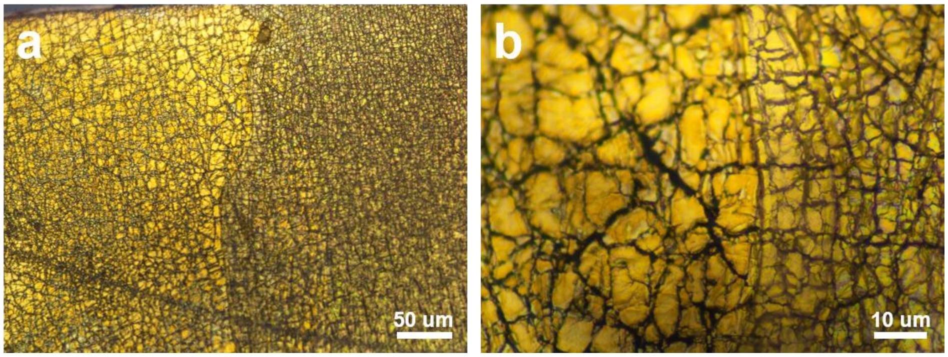


**Fig. S29.** The surface morphology of the electrode observed using an optical microscope after dissection.

**#217**

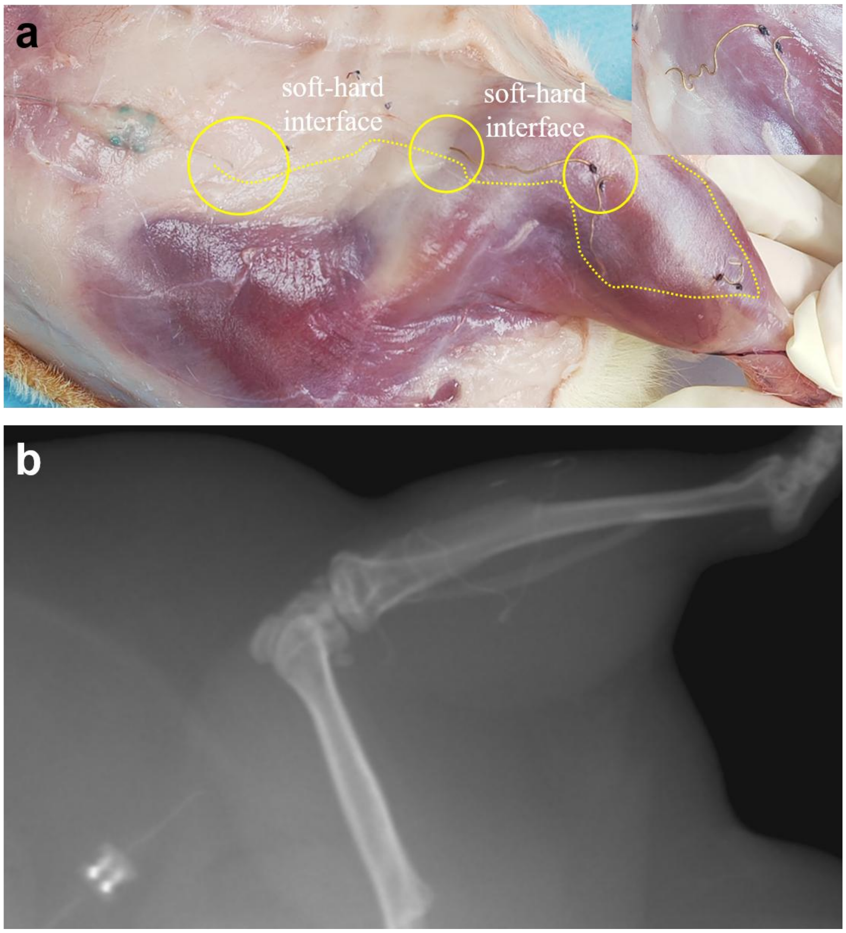


**Fig. S30.** a-b The distribution of the fiber electrodes during dissection passes through the anterior tibialis muscle, the gastrocnemius muscle, and connects to the soft-hard interface. During implantation, the electrodes are sutured to approximately secure their position in the proximal and distal ends of the anterior tibialis muscle and the fascia of the quadriceps femoris muscle they pass through. Both dissection and X-ray examination reveal that the electrodes exhibit signs of breakage at points where stress is concentrated.

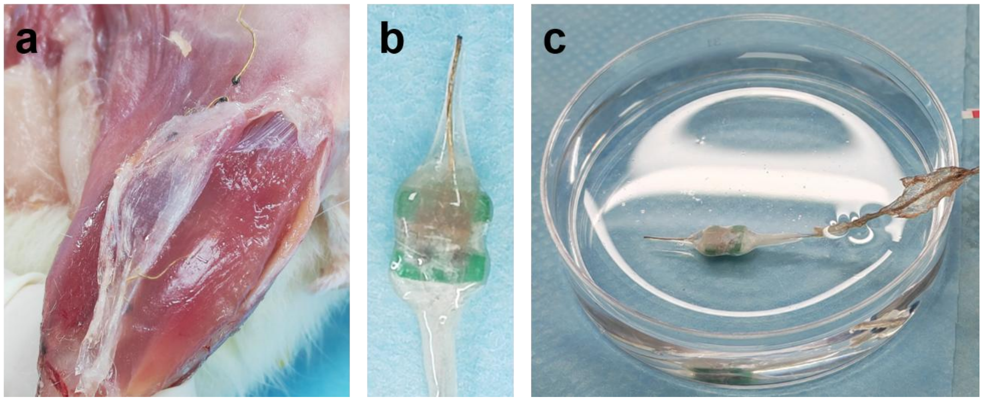


**Fig. S31.** a) The electrodes inside the muscle bundles are intact. b) The soft-hard interface is intact with no signs of water ingress, damage, or corrosion. c) Testing the seal of the interface while immersed in a PBS solution.

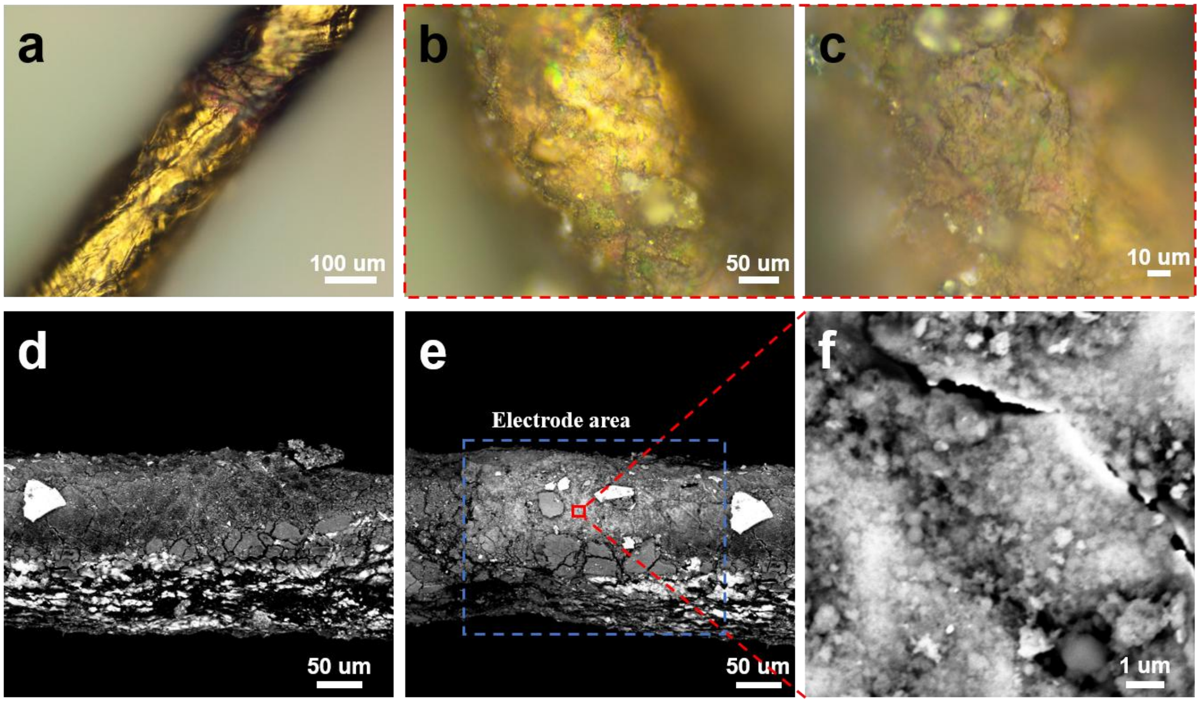


**Fig. S32.** a) The surface original fiber electrode (without implantation). b-c show the surface morphology of the electrode observed using an optical microscope after dissection, while d-f depict scanning electron microscope images.

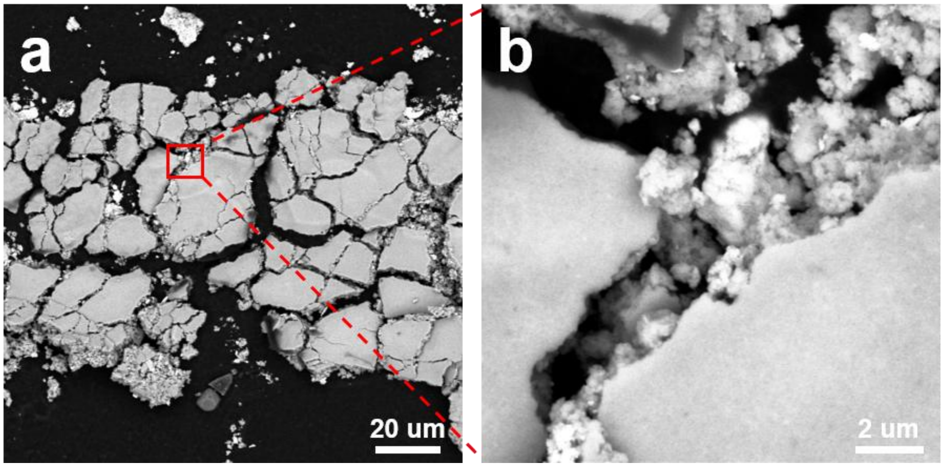


**Fig. S33.** The dissected electrode was placed on a conductive tape, and upon removing the electrode, material from the electrode surface remained on the tape. Fig. a-b show the material that adhered to the tape, as observed under an electron microscope.

**#220**

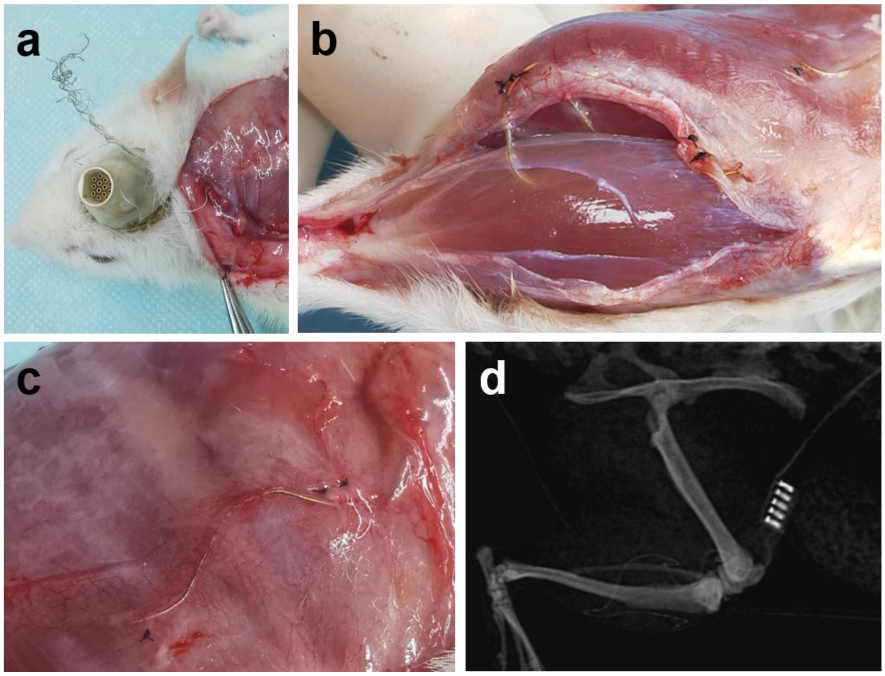


**Fig. S34.** a) The stainless steel wire at its skull interface was bent out of place (September 25, 2022). b) The fiber electrodes are distributed normally within the tibialis anterior muscle and gastrocnemius muscle, with no signs of displacement compared to the time of implantation. c-d) The soft-hard interface is intact and well-connected with the electrode, and the electrode itself remains intact.

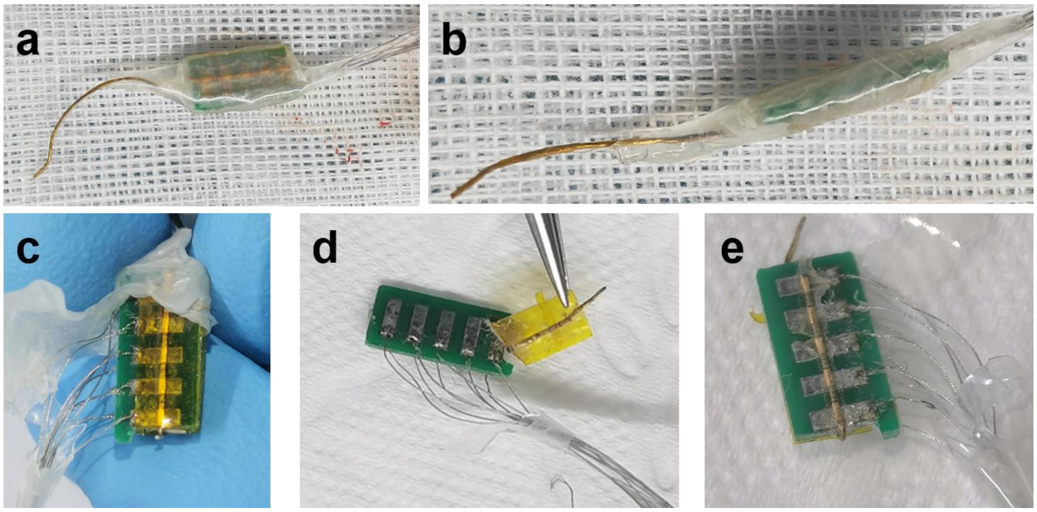


**Fig. S35.** During dissection, the soft-hard interface was observed. (a-b), and there was no internal oxidation or evidence of tissue fluid infiltration. Upon disassembling the soft-hard interface encapsulation layer (c-e), it was evident that the fiber electrode and stainless steel wire leads were securely connected to the PCB board, with no signs of corrosion.

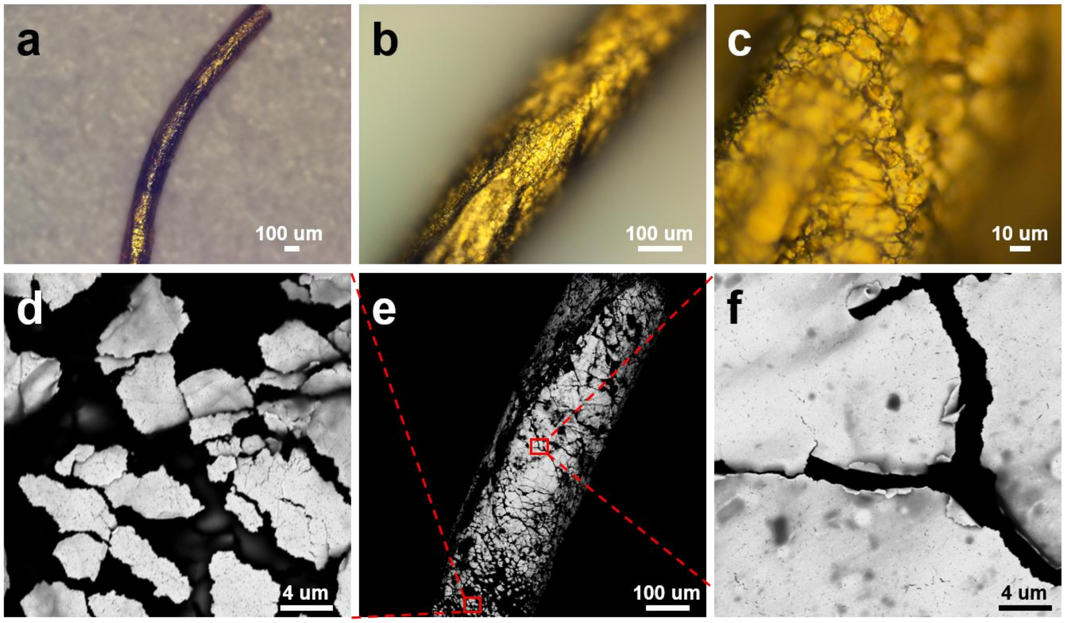

**Fig. S36.** Electrodes observed after around 43 weeks implantation. a-c) the surface morphology of the electrode observed using an optical microscope after dissection. d-f) The scanning electron microscope images.

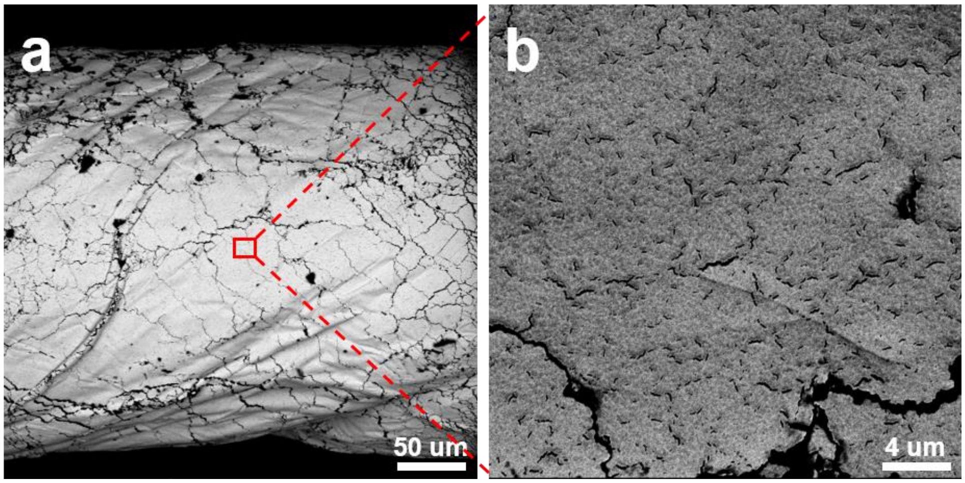

**Fig. S37.** The original surface morphology of the fiber electrode.

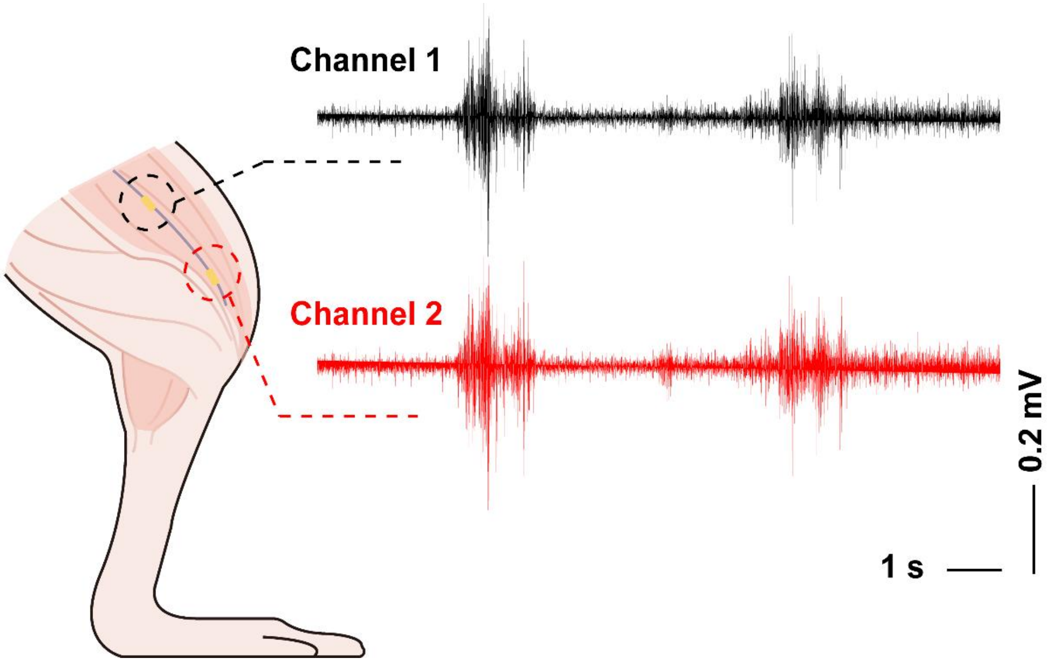


**Fig. S38.** EMG signals when the two channels of a fiber were both implanted in the anterior tibialis muscle.

The code for processing intramuscular electromyography (EMG) signals is as follows, including filtering the collected EMG signal, plotting the EMG graph, taking a screenshot of a portion of the EMG graph, and processing the central frequency value of the segmented EMG spectrogram:

```
fs = 2000;
filterRanges = [1 10; 49.5 50.5; 149.5 150.5; 249.5 250.5; 349.5 350.5; 449.5 450.5; 500 1000];
signalIndex = 27;
startpoint = fix(40.83*2e3);
endpoint = fix(42.83*2e3-1);

signal = tableData{:, signalIndex};
subsignal = tableData{startpoint:endpoint, signalIndex};

sosMatrix = [];
gainVector = [];
for i = 1:size(filterRanges, 1)
   filt_temp = designfilt('bandstopiir', 'FilterOrder', 4, 'HalfPowerFrequency1', filterRanges(i, 1),
'HalfPowerFrequency2', filterRanges(i, 2), 'SampleRate', fs);
   [num, den] = tf(filt_temp);
   [sos_temp, gain_temp] = tf2sos(num, den);
   sosMatrix = [sosMatrix; sos_temp];
   gainVector = [gainVector; gain_temp];
end
filt = dfilt.df2sos(sosMatrix, gainVector);

filteredsubSignal = filter(filt, subsignal);
filteredSignal = filter(filt, signal);

figure;
sublt = 1/fs:1/fs:size(subsignal, 1)/fs;
plot(sublt, filteredsubSignal);
axis([0 max(sublt) -2 2])
set(gca, 'XTick', 0:1:max(sublt))
set(gca, 'YTick', -2:1:2)
title(['Signal ', num2str(signalIndex)]);
ylabel('EMG signal(mv)');
xlabel('t(s)');

figure;
N = length(filteredsubSignal);
freq = (0:N/2-1) * fs / N;
spectrum = abs(fft(filteredsubSignal)/N);
plot(freq, spectrum(1:N/2));
title([' Spectrum plot ', num2str(signalIndex)]);
xlabel('Frequency (Hz)');
ylabel('Amplitude (mV)');
```

```
figure;
lt = 1/fs:1/fs:size(signal, 1)/fs;
plot(lt, filteredSignal);
axis([0 max(lt) -2 2])
set(gca, 'XTick', 0:5:max(lt))
set(gca, 'YTick', -2:1:2)
title(['Signal ', num2str(signalIndex)]);
ylabel('EMG signal(mv)');
xlabel('t(s)');

Y = fft(filteredsubSignal);
P2 = abs(Y/N);
P1 = P2(1:N/2+1);
P1(2:end-1) = 2*P1(2:end-1);
f = fs*(0:(N/2))/N;
AF_CF = sum(f'.*P1)/sum(P1);
```

**Movie S1.**
Fabrication Process of AoF-MSEA

**Movie S2.**
Implantation of wire sensor in an artificial brain under magnetic field

**Movie S3.**
Minimally-invasive implantation process of the fiber electrode

**Movie S4.**
Long-term implantable monitoring of intramuscular EMG during movement
